# Safety profiling of CAR-T cells using an organotypic human tissue platform

**DOI:** 10.64898/2026.07.25.740527

**Authors:** Machado Alice, Birsen Gary, Rajnpreht Irena, Furtado Peter Izadora, Salut Colell, Marzal Berta, Fredon Maxime, Ziegler-Martin Kai, Martinez Bedoya Darel, Fumagalli Mattia, Lene Vimeux, Davanture Suzel, Melchiore Fabien, Thomas Aurélie, Collet Mélanie, Fremand Xavier, Burroni Barbara, Laugel Bruno, Lupo-Mansuet Audrey, Prieto Mathilde, Damotte Diane, Le Doussal Jean-Marc, Luu Maik, Dutoit Valérie, Garnache-Ottou Françine, Guedan Sonia, Migliorini Denis, Donnadieu Emmanuel

**Author notes:** Correspondence to: Dr. Emmanuel Donnadieu and Dr. Alice Machado. These authors contributed equally to this work (Birsen Gary, Rajnpreht Irena).

## Abstract

CAR-T-cell-associated on-target off-tumor (OTOT) toxicity represents a major safety concern, as recognition of target antigens on healthy tissues can trigger severe and potentially life-threatening complications. Predicting OTOT toxicity remains a challenge because current preclinical models fail to capture the complexity of native human tissues. Here, we developed a human organotypic tissue platform that enables functional assessment of CAR-T-cell activity in intact human tissues across organ-specific and inflammatory contexts. Using a panel of clinically relevant CAR-T-cell products with known OTOT toxicities, we demonstrate that the platform faithfully recapitulates clinically observed tissue-specific toxicity profiles. CAR-T cells targeting EGFR, HER2, and mesothelin induced inflammatory and cytotoxic responses in healthy human lung tissue, whereas CD19 CAR-T cells remained inactive. We further show that OTOT toxicity cannot be reliably predicted from antigen abundance alone but instead results from the integration of multiple target-dependent determinants, including CAR affinity, inflammatory context, antigen accessibility, and effector-cell dose. The platform also enables quantitative assessment of inflammatory and cytotoxic responses and supports evaluation of pharmacological and CAR design-based strategies to mitigate toxicity. Together, this work establishes the first human organotypic platform for functional modeling of CAR-T-cell-associated OTOT toxicity, providing a clinically relevant framework for preclinical safety evaluation and the rational development of safer engineered cell therapies.

**Graphical abstract:** 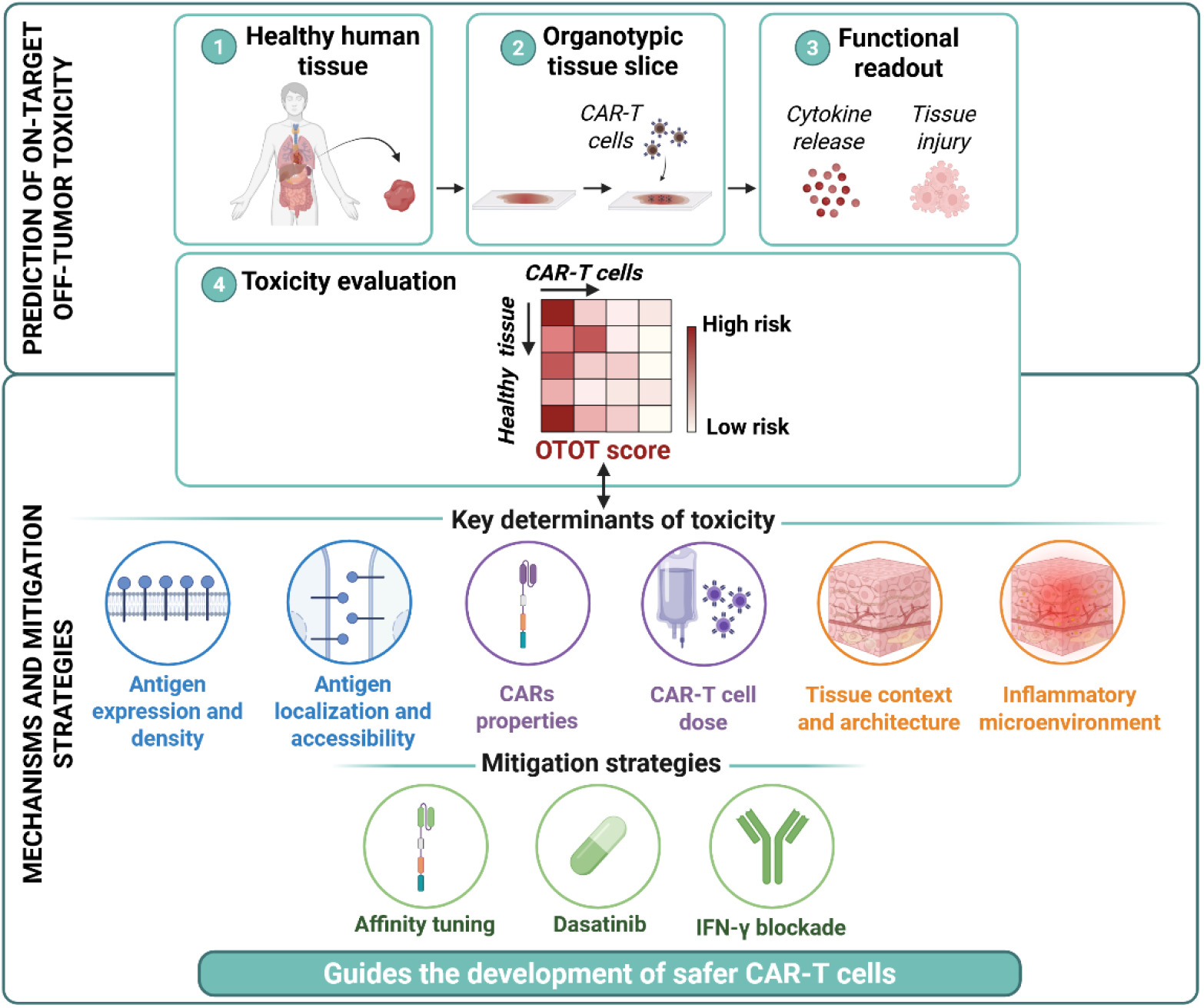

## Introduction

Chimeric antigen receptor T-cell (CAR-T) therapy has emerged at the forefront of immuno-oncology research and has shown significant clinical success in hematologic malignancies, with seven CAR-T cell therapies currently approved. However, their use is associated with significant and diverse toxicities ^1–3^.

Adverse events arise from multiple sources, including treatment-related toxicities independent of CAR-T cells (e.g., lymphodepleting conditioning), systemic immune activation-such as cytokine release syndrome (CRS), immune effector cell-associated neurotoxicity syndrome (ICANS), and immune effector cell-associated hemophagocytic syndrome (IEC-HS)-as well as antigen-related toxicities, notably on-target off-tumor (OTOT) effects ^3–5^. OTOT toxicity results from CAR-T cell recognition of shared tumor-associated antigens (TAAs) on normal tissues, leading to direct CAR-T-mediated cytotoxicity against healthy cells. For example, CD19-targeted CAR-T cells induce predictable lineage-specific B-cell aplasia, which can be effectively managed ^6^.

CAR-T therapy is not yet approved for solid tumors, largely due to the lack of tumor-specific antigens, increasing OTOT risk. In most cases, TAAs are also expressed on normal tissues, making toxicity less predictable and more difficult to control ^7^.

CAR-T cell-associated toxicities may affect multiple organs. Pulmonary involvement during CRS is severe and can progress to acute respiratory distress syndrome (ARDS). Moreover, OTOT toxicity affecting the lung has been associated with life-threatening complications in several studies ^8–12^. In addition, CAR-T cells transiently accumulate in the pulmonary circulation after infusion, increasing antigen exposure in this organ ^13,14^. Additional factors, such as CAR design and the administered cell dose, influence activation thresholds, reactivity toward normal tissue and toxicity severity ^15–19^. This risk is further modulated by the state of normal tissue, which can be reshaped by the tumor microenvironment and prior treatments, leading to local inflammatory changes.

This complexity underscores the need for preclinical models that integrate the multiple parameters governing CAR-T cell toxicity in normal tissues. Current OTOT risk assessment still largely relies on target antigen detection in healthy tissues. Although essential, this approach has important limitations. Immunohistochemistry depends on antibody quality, staining conditions and detection thresholds, while transcriptomic analyses do not necessarily reflect protein abundance, localization or accessibility. Moreover, antigen expression alone is not sufficient to predict toxicity, as CAR-T cell activation is also shaped by antigen density, spatial distribution, epitope accessibility, receptor affinity, effector dose and the local tissue environment. Inflammatory or stress signals within normal tissues may further modulate both antigen expression and tissue sensitivity to CAR-T cell-mediated injury ^20,21^. These determinants are poorly captured by conventional 2D cultures, xenograft models or expression-based screening strategies. Organoids partially reconstruct tissue architecture and heterogeneity, but still lack the full cellular and stromal complexity of intact tissues ^22,23^. We therefore established an *ex vivo* organotypic model using fresh human tissue slices, preserving native architecture, cellular diversity and key biological properties, including antigen expression, cytokine accumulation and cellular responses to external stimuli ^24–26^.

In this study, we used a diverse CAR-T panel to evaluate the platform across functional and safety-relevant contexts. Initial experiments with EGFR-, HER2-, and mesothelin-(MSLN) CAR-T cells associated with OTOT toxicities-alongside-CD19 CAR-T cells as a non-toxic control, validated the system through tissue-specific cytokine release and cytotoxicity ^9–11^. Predictive performance was further supported by ROR1 CAR-T cells in murine models, where dose-dependent pulmonary toxicity was recapitulated *in vivo*. Application across multiple healthy cynomolgus macaque tissues with glioblastoma-targeting CAR-T cells demonstrated the platform’s suitability for multi-organ screening. The system also identified IFN-γ signalling as a driver of CD123 CAR-T-mediated activation and enabled mitigation strategies, including anti-IFNγ antibodies, dasatinib, and CAR affinity tuning.

Importantly, our findings demonstrate that OTOT toxicity is highly target-dependent and cannot be predicted solely by the presence of a target antigen. Rather, toxicity emerges from the specific CAR-target interaction, integrating target expression, antigen density and accessibility, CAR affinity, effector-cell dose, tissue susceptibility, and local inflammatory signals. Consequently, each target antigen may generate a distinct toxicity profile and functional outcome upon CAR-T cell engagement, highlighting the need for functional models that evaluate CAR-T cell reactivity within native tissue environments. By preserving human tissue architecture and biological responses, our approach enables direct assessment of CAR-T cell activity against physiological targets and provides a framework to identify toxicity mechanisms and develop mitigation strategies before clinical translation.

## Results

### Organotypic lung slices recapitulate the clinical OTOT toxicity of EGFR, HER2, and Mesothelin CAR-T cells

To predict OTOT toxicities, we used an organotypic platform based on fresh healthy human lung tissue collected from areas distant from the tumor within lobectomy specimens, co-cultured with T cells (**Fig. 1A**). We first characterized lung slices using spatial transcriptomics, as strong intrinsic autofluorescence limits fluorescence-based microscopy. A CAR-specific probe enabled selective detection of engineered T cells, with signal observed only in the HER2 CAR-T condition, confirming assay specificity and demonstrating CAR-T cell infiltration into the tissue (**Fig. Sup 1A, 1B**). The analysis further revealed the persistence of the major pulmonary cell populations, including endothelial, epithelial, fibroblast, and smooth muscle cells, as well as immune populations such as T and B lymphocytes, monocytes, and macrophages. These populations remained organized in spatially coherent structures, indicating preservation of tissue architecture and cellular diversity during culture (**Fig. Sup 1C**). We then tested EGFR, HER2 high-affinity and MSLN CAR-T cells, which have been associated with severe OTOT pulmonary toxicities in clinical trials ^9–11^. As a negative control, we used CD19-targeting CAR-T cells, which are not expected to induce OTOT pulmonary toxicity. Given the variability in CAR transduction efficiency across constructs (**Fig. Sup. 2A**), CAR-T cell responses were analyzed at a standardized 30% CAR-positive fraction among total T cells. Following the conventional approach used for OTOT risk assessment, we first evaluated target antigen expression in healthy lung tissue by immunohistochemistry (IHC). CD19, EGFR, HER2, and MSLN were detected at low levels with inter-patient variability (**Fig. 1B**). Despite this overall low antigen detection, EGFR-, HER2-, and MSLN-targeting CAR-T cells were selected because they have been associated with clinically reported pulmonary OTOT toxicities. We therefore proceeded to evaluate CAR-T cell activity using our *ex vivo* human lung tissue model.

**Fig. 1:**
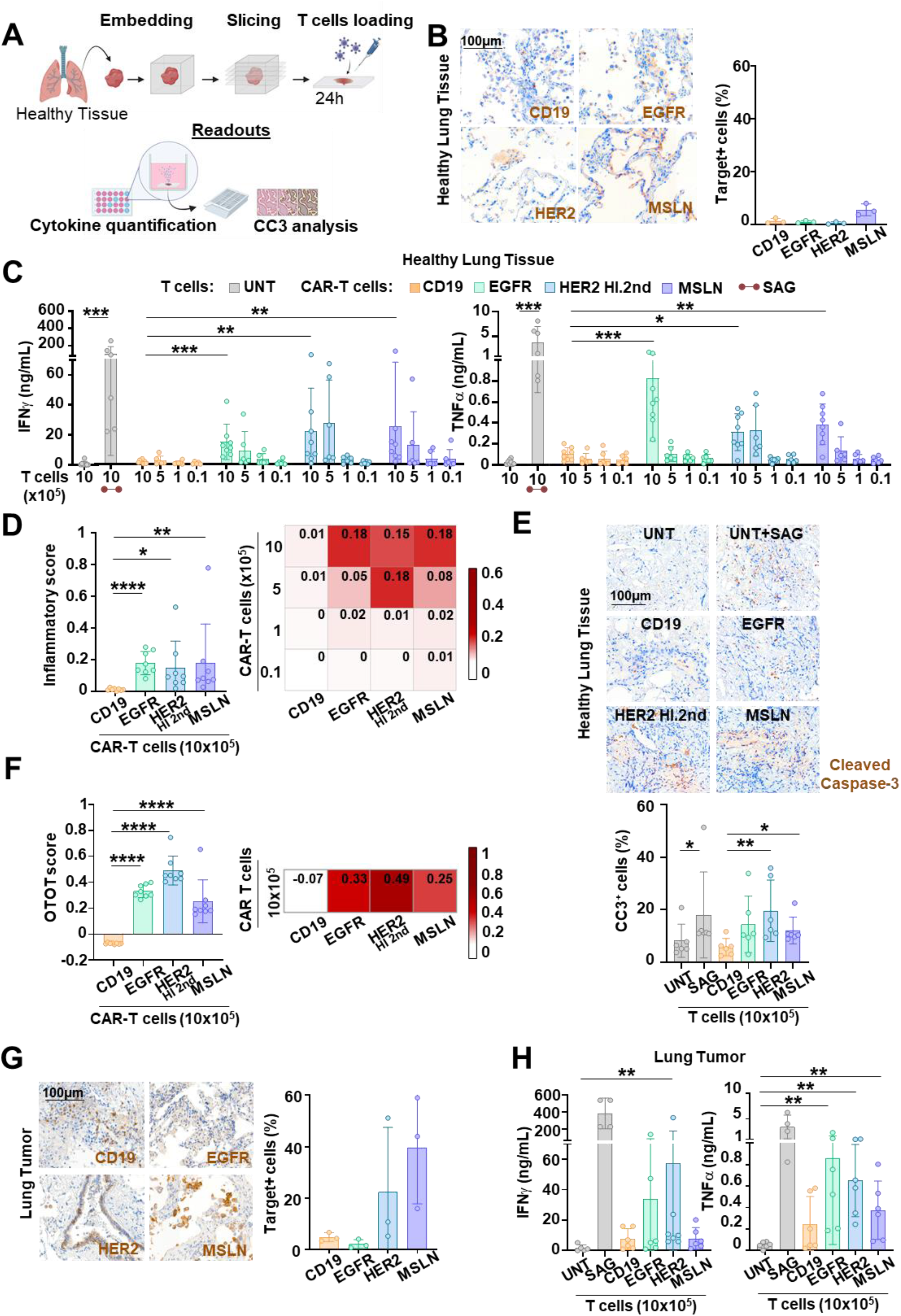
On-target off-tumor toxicity is observed with EGFR, HER2, and MSLN CAR-T cells but not with CD19 CAR-T cells, consistent with clinical data. **A)** Schematic of the organotypic slice model. Healthy tissues were harvested, sliced, and co-cultured with CAR-T cells. Each slice received 1×10⁶ total T cells at different CAR-T/UNT ratios (10/0, 5/5, 1/9, and 0.1/9.9). After 24 h, cytokine secretion and cleaved caspase-3 (CC3) expression were assessed. **B)** IHC analysis and quantification of CD19, EGFR, HER2, and MSLN expression in healthy lung tissue (mean ± SD, n=4). **C)** IFN-γ and TNF-α secretion following co-culture of healthy lung slices with CD19, HER2 high-affinity, EGFR, and MSLN CAR-T cells. UNT and SAG-treated UNT served as negative and positive controls, respectively (n=3 tissue donors; n=3 T-cell donors; duplicate/triplicate slices) **D)** Inflammatory score calculated from IFN-γ and TNF-α secretion. Bar graph shows 10×10⁵ CAR-T cells per slice; heatmap summarizes all tested doses. Scores normalized from 0 (UNT) to 1 (SAG). **E)** Images showing CC3 (brown) detected by IHC. Quantification of CC3+ cells is shown with UNT and CAR-T cells at 10×10^5^ cells per slice, n = 6 sections. **F)** OTOT score calculated from IFN-γ and TNF-α concentrations and the % of CC3+ cells. Bar graph shows 10×10⁵ CAR-T cells per slice (means ± SD); Heatmap summarizes all tested doses. Scores were normalized between 0 (UNT) and 1 (SAG) **G)** IHC staining and quantification of CD19, EGFR, HER2 and MSLN expression in lung adenocarcinoma slices (means ± SD). **H)** IFN-γ and TNF-α secretion following co-culture of lung adenocarcinoma slices with CAR-T cells (10×10⁵ CAR-T cells per slice). UNT and SAG-treated UNT served as controls (n = 3 tissue donors; n = 3 T-cell donors; duplicate slices per condition). Statistical significance was determined using a two-tailed Mann-Whitney test or one-way ANOVA with Tukey’s multiple-comparison test. *P < 0.05; **P < 0.01; ***P < 0.001; ****P < 0.0001.

CAR-T cells targeting HER2, EGFR, and MSLN secreted high levels of the pro-inflammatory cytokines IFN-γ and TNF-α in a dose-dependent manner following co-culture with healthy lung tissue slices, whereas CD19 CAR-T and UNT cells did not induce cytokine release. As a positive control, superantigen (SAG) stimulation induced robust cytokine secretion, confirming tissue responsiveness in the assay (**Fig. 1C**). To provide a quantitative overview of inflammatory responses, we calculated an inflammation score based on combined IFN-γ and TNF-α levels, normalized to UNT cells (negative control, score = 0) and SAG treatment (positive control, score = 1). The score reflected CAR-T cell-induced cytokine responses, with EGFR-, HER2-, and MSLN-targeting cells displaying elevated inflammatory scores, while CD19 CAR-T cells remained at baseline (**Fig. 1D**). Apoptotic responses were then assessed via cleaved caspase-3 (CC3) staining on the slices. HER2-EGFR-MSLN-targeting CAR-T cells induced CC3 expression whereas no increase was observed with CD19 CAR-T or UNT cells (**Fig. 1E**). An integrated OTOT score combining inflammation and CC3 (UNT (negative control, score = 0) and normalized to UNT and SAG controls revealed significantly higher toxicity for EGFR, HER2, and MSLN CAR-T cells compared to CD19 (**Fig. 1F**).

In the absence of target cells, CAR-T cells did not secrete cytokines, whereas co-culture with antigen-positive tumor cells induced strong cytokine responses (**Fig. Sup. 2B**).

Overall, although CD19, EGFR, HER2, and MSLN showed low antigen detection in healthy lung tissue by IHC, only EGFR-, HER2-, and MSLN-targeting CAR-T cells induced inflammatory and apoptotic responses. These results show that the organotypic lung slice assay functionally discriminates CAR-T cells associated with clinical pulmonary OTOT toxicity from a non-toxic CD19 control.

### CAR-T cell responses in lung adenocarcinoma suggest microenvironmental modulation of CAR-T activation

We next evaluated CAR-T cell activity in lung adenocarcinoma samples, enabling comparison of tumor and non-tumor lung tissue contexts from the same samples. Tumor antigen expression was heterogeneous across CD19, EGFR, HER2, and MSLN, reflecting their variable prevalence in lung adenocarcinoma (**Fig. 1G**). Compared with non-tumoral lung tissue, CD19 and EGFR expression showed only modest changes in tumor samples, and this was associated with similar cytokine responses between the two tissue contexts. In contrast, HER2 and MSLN expression were increased in some tumor samples, but this did not translate into higher CAR-T cell cytokine secretion compared with non-tumoral lung tissue. This was particularly evident for MSLN-targeting CAR-T cells, which showed limited activation in tumor slices despite detectable target enrichment (**Fig. 1H**).

Thus, within intact lung tissues, CAR-T cell activation was not simply proportional to antigen expression as assessed by IHC. These results suggest that additional parameters, such as antigen accessibility, spatial distribution, local tissue organization and tumor-associated microenvironmental constraints, may influence CAR-T cell responses. The tissue slice model preserves key features of the native tumor microenvironment that continue to modulate CAR-T cell activity *ex vivo*.

### Organotypic lung slices reliably recapitulate antigen-specific CAR-T cell activation and cytotoxicity

To confirm that organotypic lung slices capture CAR-T cell activation within the tissue environment, we performed spatial transcriptomics on HER2 CAR-T cell-treated slices. IHC was used to delineate HER2+ regions, providing spatial annotation and validation for the transcriptomic analysis. CAR-T cells were frequently detected in in spatial proximity to these regions, where they appeared clustered and exhibited expression of activation-associated genes. In these areas, we also observed increased expression of apoptotic markers in surrounding cells, consistent with localized CAR-T-associated cytotoxic activity (**Fig. 2A**).

**Fig. 2:**
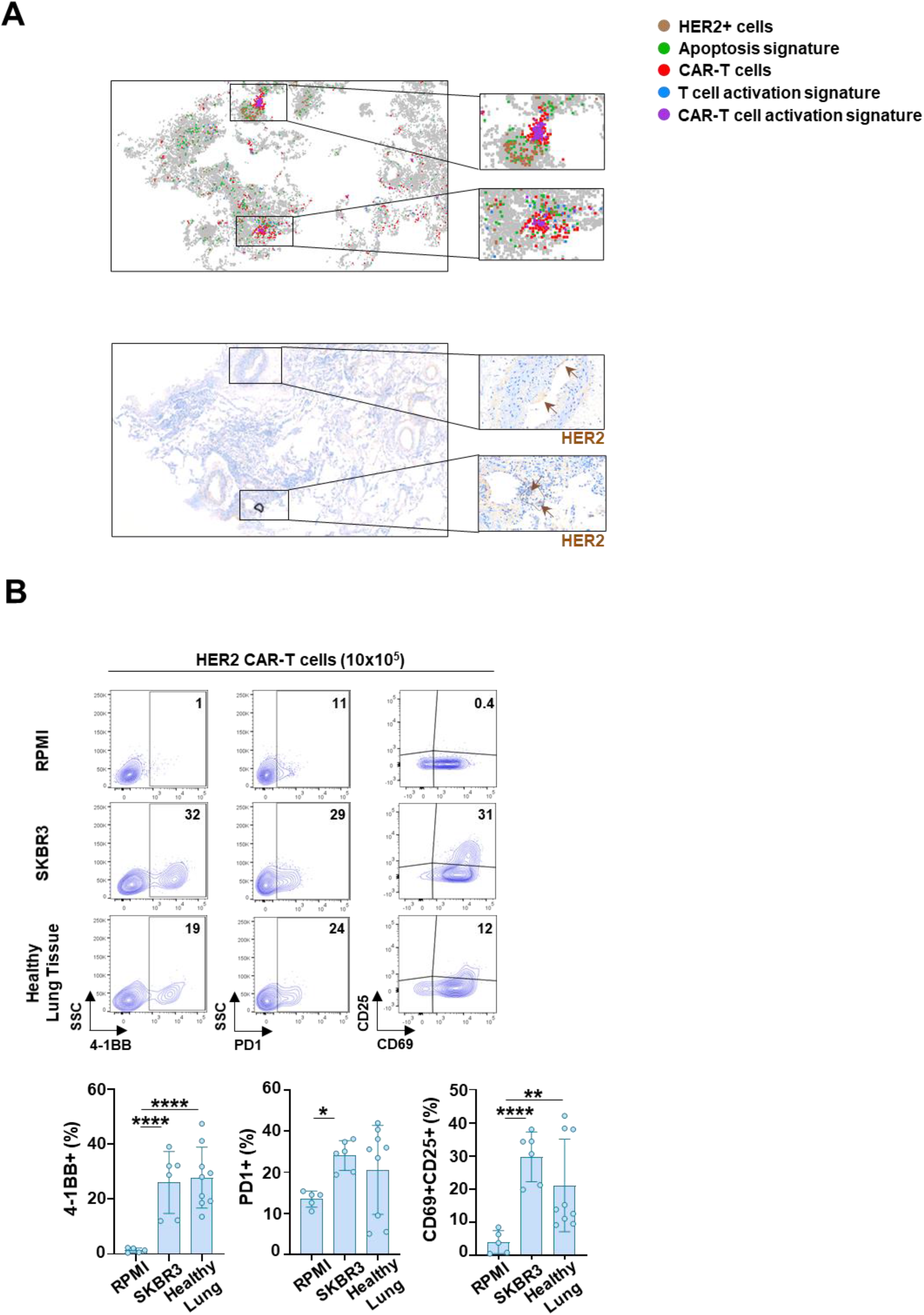
Organotypic lung slices recapitulate antigen-specific HER2 CAR-T activation and cytotoxicity. A) Spatial transcriptomic analysis of HER2 CAR-T-treated organotypic lung slices. Map showing CAR-T cell activation and apoptosis signals across the tissue. Each dot represents a capture spot, with signal intensity reflecting CAR-T activation and apoptotic marker expression (top). HER2 expression assessed by IHC (brown and arrow) (bottom). **B)** Dot plots showing activation marker expression (4-1BB, PD-1, CD69, and CD25) in HER2 CAR-T cells after 24 hours of co-culture with healthy lung tissue in the presence of RPMI control or SKBR3 target cells. Corresponding quantification is shown below. Statistical significance was determined using a two-tailed unpaired t-test; *, p < 0.05; **, p < 0.01; ***, p < 0.001; **, p < 0.0001.

We confirmed that HER2 CAR-T cells upregulated early activation markers, including 4-1BB, PD-1, CD69 and CD25, after 24 hours of co-culture with healthy lung tissue. Importantly, this activation profile was consistent with cytokine secretion patterns, supporting that cytokine readouts accurately reflect antigen-specific CAR-T cell activation (**Fig. 2B**).

### Organotypic lung slices mirror murine pulmonary OTOT toxicity of ROR1 CAR-T cells

We next sought to evaluate whether our *ex vivo* organotypic assay could accurately reflect in vivo responses and model pulmonary toxicity. To this end, we used ROR1-targeting CAR-T cells, with approximately 30% CAR-positive cells, which are known to induce pulmonary OTOT toxicity in mice^13,27^. We confirmed the absence of basal cytokine secretion under control conditions and demonstrated robust cytotoxic activity of ROR1 CAR-T cells against a ROR1-expressing cell line (**Fig. Sup. 3A, 3B**).

As a first step, we established the *in vivo* model by injecting murine ROR1 CAR-T cells (1 or 10 × 10⁶) or UNT T cells (10 × 10⁶) into CD45.2+ C57BL/6 mice bearing subcutaneous pancreatic (PAN02) tumors. Three days after injection, both UNT and CAR-T cells were detected in the healthy mouse lungs. Immunofluorescence revealed that ROR1 CAR-T cell presence was dose-dependent (**Fig. 3A**). Moreover, CAR-T cells preferentially accumulated in ROR1-expressing lung regions, indicating specific targeting (**Fig. 3B**).

**Fig. 3:**
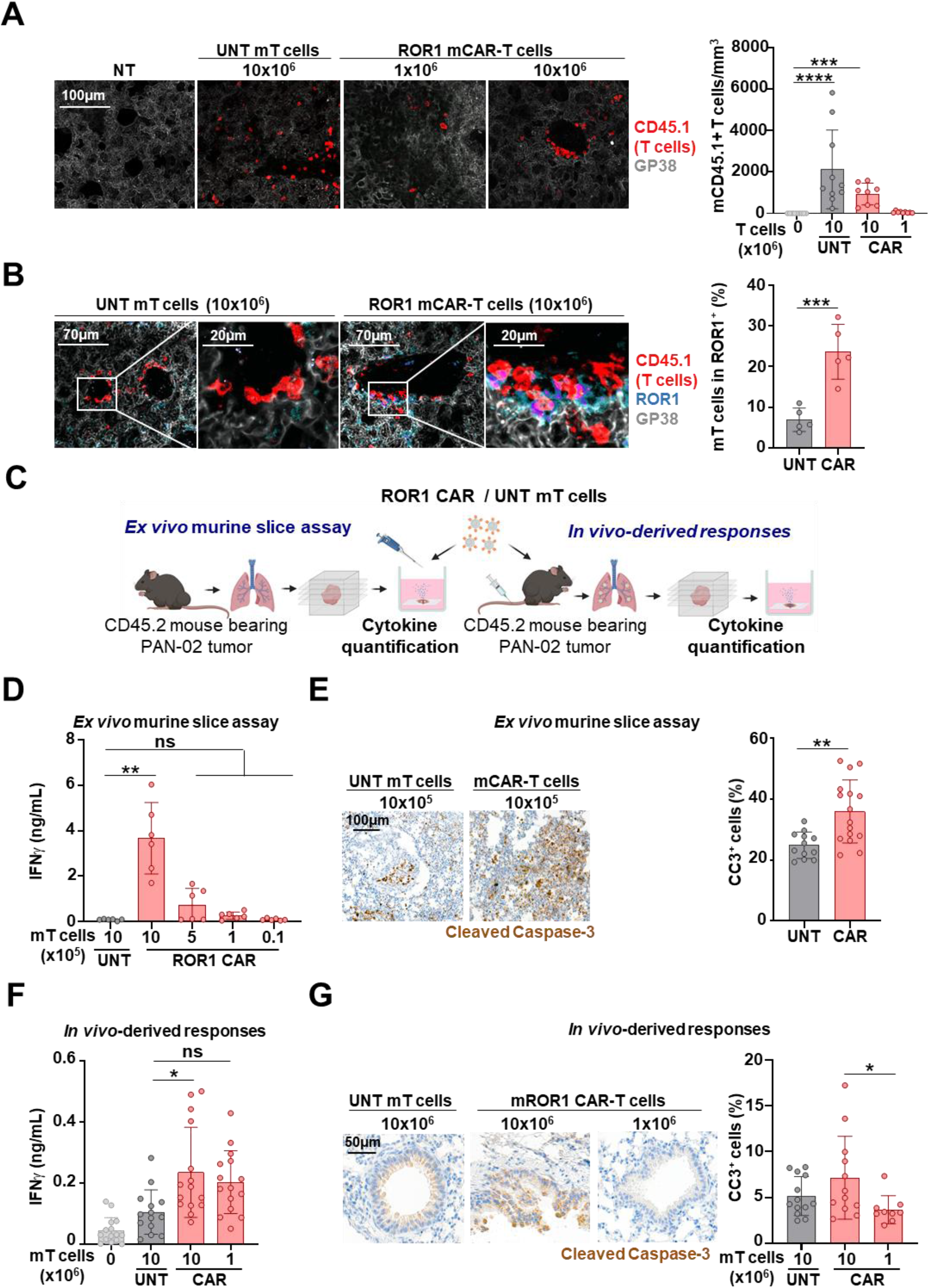
*Ex vivo* organotypic model faithfully recapitulates *in* vivo ROR1 murine CAR-T cell-induced pulmonary OTOT **A)** Confocal images of lung sections stained for CD45.1 (CAR-T cells, red) and GP38 (gray) from tumor-bearing mice injected intravenously with ROR1 CAR-T cells (10×10⁶ or 1×10⁶), UNT (10×10⁶), or non-injected controls. Bar graph shows CD45.1⁺ cell quantification (n = 5). **B)** Representative lung images from mice injected with 10×10⁶ CAR-T or UNT murine cells, stained for CD45.1 (red, T cells), ROR1 (blue), and GP38 (gray). Quantification of CD45.1⁺ cells within the ROR1⁺ compartment is shown (n = 3, duplicate). **C)** Experimental design; Lungs from tumor-bearing mice and injected with ROR1 murine CAR-T cells (10×10⁶ or 1×10⁶) or UNT (10×10⁶) were sliced and cultured for 24 hours before collecting supernatants (right). Healthy lungs from tumor-bearing mice were sliced and cultured with CAR-T or UNT cells for 24 hours, then supernatants were collected (left). Cytokine levels were measured. **D)** Bar graph showing mean IFN-γ secretion by CAR-T cells added at different doses in the *ex vivo* organotypic model (means ± SD; n = 3 in duplicate). **E)** IHC images showing CC3 positive cells (brown) from lung slices treated with UNT and CAR-T cells (1×10^6^) in the *ex vivo* organotypic model (n = 12-16 sections per group) **F)** Quantification of IFN-γ in supernatants from lung slices of mice injected intravenously with murine CAR-T cells (10×10⁶ or 1×10⁶), UNT cells (10×10⁶), or non-treated mice, presented as mean ± SD (n = 5 per group, triplicates). **G)** IHC images of lung slices from mice injected intravenously with UNT cells or CAR-T cells, stained for CC3 (brown) (mean ± SD; n = 8-11 sections). Statistical significance was determined using Kruskal-Wallis and Mann-Whitney tests; *P < 0.05, **P < 0.01, ***P < 0.001, ****P < 0.0001.

We then used the murine model to assess the relevance of the *ex vivo* organotypic assay by directly comparing CAR-T cell responses across both systems. Healthy mouse lungs collected three days post-injection, confirmed to contain either CAR-T or UNT cells (**Fig. 3A**), were sliced and cultured *ex vivo* for 24 hours. In parallel, we performed the standard organotypic assay by applying the same ROR1 CAR-T and UNT cells to healthy lung slices obtained from untreated mice (**Fig. 3C**). As expected, both models showed increases in IFN-γ secretion. Specifically, IFN-γ and CC3 levels were significantly elevated at a dose of 1×10⁶ CAR-T cells in the organotypic assay, and at 10×10⁶ cells in the in vivo setting (**Fig. 3D, 3E, 3F, 3G**). Notably, absolute cytokine levels differed between *ex vivo* and *in vivo* settings, most likely due to differences in sampling: in *ex vivo* slices, IFN-γ accumulates in the supernatant from the time of CAR-T cell addition, whereas *in vivo* it is assessed in lung tissue four days post-CAR-T infusion, capturing only a transient release. Altogether, these results show functional concordance between the *in vivo* and *ex vivo* organotypic systems, supporting the ability of the *ex vivo* assay to recapitulate CAR-T cell-mediated toxicity. Both models consistently highlight a dose-dependent pulmonary response of ROR1 CAR-T cells, underscoring the importance of CAR-T cell dose in pulmonary OTOT toxicity assessment.

### Organotypic lung slices serve as a platform to pre-screen CAR-T cell OTOT toxicity

We expanded the platform to assess OTOT toxicity across multiple tissues and evaluate its potential as a pre-screening tool for CAR-T cell safety. We tested human mRNA CAR-T cells targeting glioblastoma-associated antigens (BCAN, CSPG4, PTPRZ1, and TNC ^28–30)^ on organotypic slices from cynomolgus macaque tissues, including cerebellum, cortex, peripheral nerve, heart, lung, liver, skin, colon, pancreas, and kidney (**Fig. 4A, Fig. Sup. 4A**). After 24 hours of co-culture, most CAR-T cell constructs showed minimal activity in healthy tissues, except for TNC CAR-T cells (**Fig. 4B; Fig. Sup. 4B**). Calculation of the inflammatory score confirmed that TNC CAR-T cells induced strong pro-inflammatory activity across multiple organs, comparable to the positive control (SAG) (**Fig. 4C**). Consistent with the correlation between cytokine release and cytotoxicity, elevated cytokine production in the TNC CAR-T cell conditions was associated with CC3 detection in liver slices (**Fig. 4D**). Basal cytokine secretion was absent, confirming assay specificity, and CAR-T cell functionality was validated using antigen-expressing cell lines (**Fig. Sup. 4C**).

**Fig. 4:**
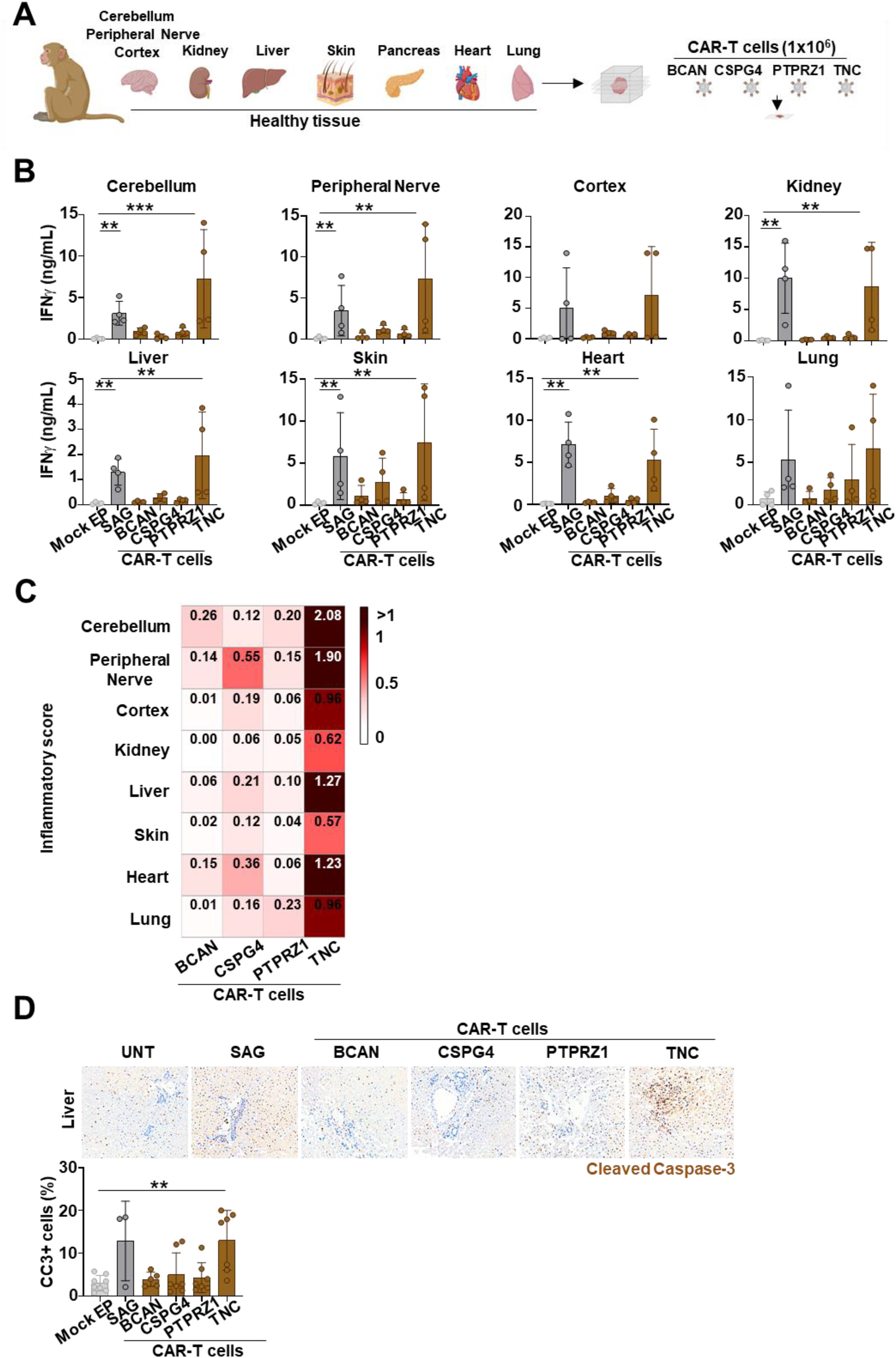
Organotypic slices from cynomolgus macaque tissues reveal broad OTOT toxicity potential of TNC-targeting CAR-T cells. **A)** Schematic of the organotypic assay showing glioblastoma-targeting CAR-T cells (BCAN, CSPG4, PTPRZ1, and TNC) tested on healthy cynomolgus macaque tissue slices, including cerebellum, cortex, peripheral nerve, kidney, liver, skin, heart, pancreas, and lung. **B)** Bar graph showing IFN-γ secretion by the different mRNA CAR-T cells and mock electroporated (mock EP) T cells (1×x10^6^ cells/slice) across various healthy tissues. Positive control: SAG and mock EP T cells (SAG). Data represent n = 2 cynomolgus macaques, performed in duplicate (mean ± SD). **C)** Heatmap displaying the inflammatory score derived from IFN-γ and TNF-α levels. Values were normalized on a scale from 0 (baseline response from mock EP T cells cells) to 1 (maximum response induced by SAG), enabling standardized comparison across all conditions. **D)** IHC images of liver slices treated with the different CAR-T cells (1×x10^6^ cells), stained for CC3 (brown). The percentage of CC3+ cells is shown (mean ± SD), based on n = 7 sections per condition. Statistical analyses were performed using the Kruskal-Wallis test followed by Mann-Whitney comparisons. Significance levels were indicated as follows; *, p < 0.05; **, p < 0.01; ***, p < 0.001; ****, p < 0.0001.

TNC CAR-T cells were the only construct to induce robust inflammatory responses across multiple healthy tissues, highlighting a potential OTOT liability that was readily detected by the platform. While the precise mechanism underlying this reactivity remains to be determined, it may reflect recognition of physiological TNC expression in healthy tissues, tissue-processing-associated changes in antigen accessibility ^31^, or species-specific cross-reactivity with cynomolgus macaque proteins.

Collectively, these findings demonstrate that the organotypic tissue platform can identify potential activation across a broad range of organs. This versatility supports its use as a preclinical platform for safety assessment and therapeutic target prioritization prior to clinical development.

### Organotypic lung slices reveal IFN-γ-dependent modulation of CAR-T cell reactivity

Following validation of the platform’s robustness, we next investigated its potential to go beyond OTOT toxicity prediction and capture dynamic changes in the tissue microenvironment. In particular, we focused on inflammatory signaling, with a specific emphasis on IFN-γ, a key cytokine involved in microenvironmental reprogramming. To assess whether organotypic lung slices can capture such environment-driven changes ^34–36^, we pretreated the slices with recombinant IFN-γ (rIFN-γ) prior to CAR-T cell exposure. CD123 was chosen as a CAR-T target of particular interest, given its upregulation on endothelial cells under inflammatory conditions and its known involvement in CAR-T cell-related toxicities^35,36^. IFN-γ preconditioning enhanced CD123 expression and selectively increased CD123 CAR-T cell activation, as reflected by elevated inflammatory cytokine secretion, whereas no comparable effect was observed for CD19-, EGFR-, HER2-, or MSLN-targeted CAR-T cells (**Fig. Sup. 5A**; **Fig. 5A, 5B, 5C)**. These results suggest that IFN-γ selectively modulates CAR-T responses in a target-dependent manner, highlighting the ability of the platform to capture microenvironment-driven changes in tissue reactivity.

**Fig. 5:**
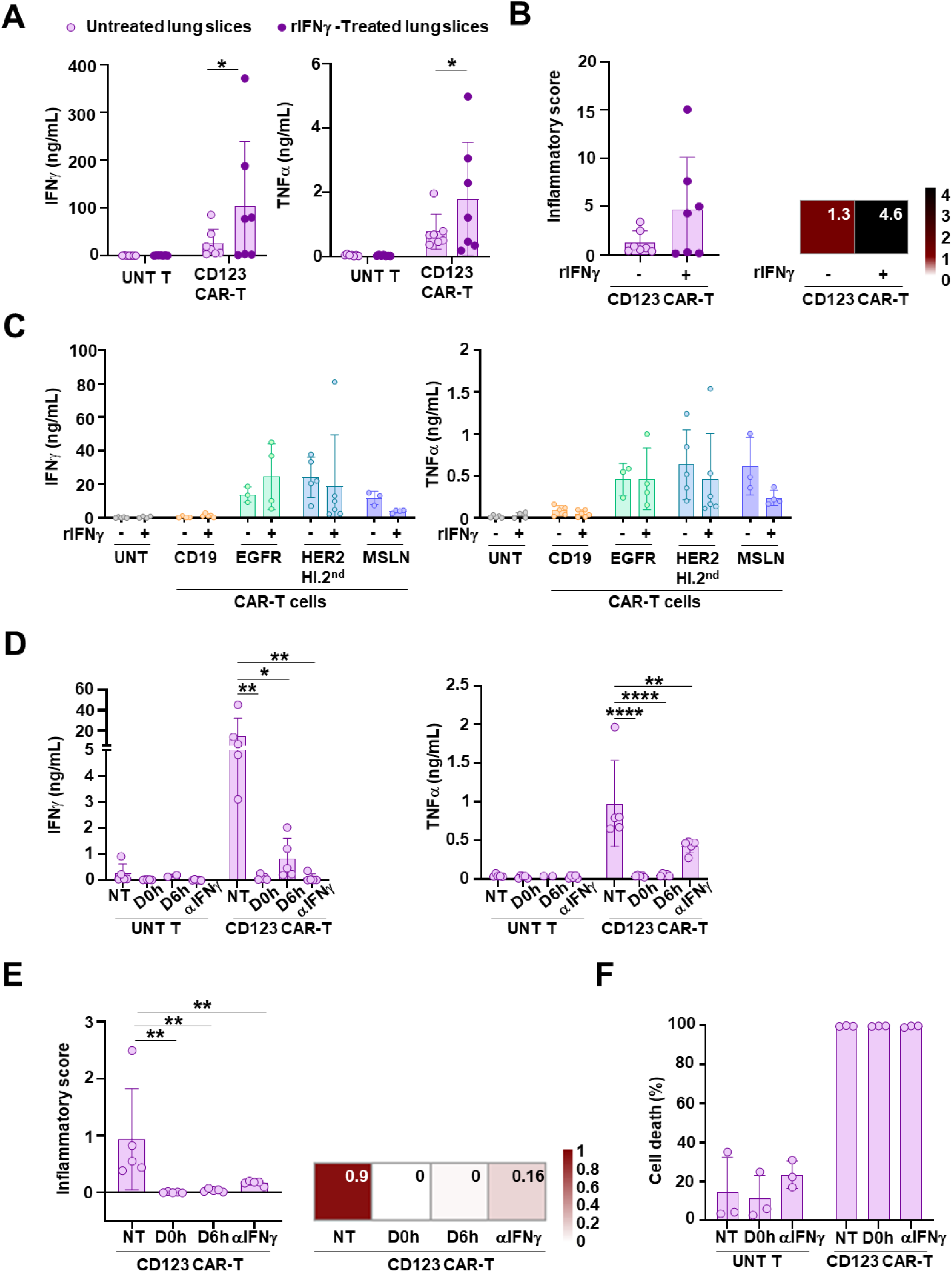
Organotypic lung slices reveal IFN-γ-dependent modulation of CD123 CAR-T-associated pulmonary OTOT toxicity and its pharmacological mitigation. **A)** IFN-γ and TNF-α secretion by CD123 CAR or UNT-T cells (10×10⁵ cells per slice) following 24 h co-culture with untreated or rIFN-γ-pretreated (10 pg/mL) lung slices (n = 3 tissue donors; n = 3 T-cell donors; duplicate slices) **B)** Inflammatory score calculated from IFN-γ and TNF-α secretion. Bar graph shows normalized scores for 10×10⁵ cells per slice; heatmap summarizes mean values across conditions. Scores were normalized on a scale from 0 (UNT) to 1 (SAG). **C)** IFN-γ and TNF-α secretion by CD19, EGFR, HER2 high-affinity and MSLN CAR or UNT-T cells (10×x10⁵ cells per slice) following co-culture with untreated or rIFN-γ-pretreated lung slices (10 pg/mL) (n = 2 tissue donors). **D)** IFN-γ and TNF-α secretion by CD123 CAR-T cells under untreated conditions or following dasatinib treatment (50 nM, added at 0 or 6 h) or anti-IFN-γ antibody treatment (10 µg/mL) (n = 2, duplicates or triplicates). **E)** Normalized inflammatory scores for the conditions shown in D, presented as bar graphs and heatmaps. **F)** Bar graphs showing CD123 CAR-T-mediated cytotoxicity against CAL-1 target cells (E:T ratio = 1:1, 24h) in the presence or absence of anti-IFN-γ (10 µg/mL) or dasatinib (50 nM). UNT T cells were used as control. Data represent mean ± SD (n = 3). Statistical significance was determined using a two-tailed Mann-Whitney test or a one-way ANOVA with Tukey’s multiple comparison test; *, p<0.05; **, p<0.01; ***, p<0.001; **\*\*\*\***, p<0.0001.

### Pharmacologic strategies decrease pulmonary OTOT toxicity of CD123 CAR-T cells

Given the prominent role of IFN-γ in shaping CAR-T cell-associated OTOT responses, and the robust IFN-γ secretion observed in CD123 CAR-T conditions, we next assessed whether blocking IFN-γ could reduce OTOT toxicity. Treatment with the anti-IFN-γ antibody resulted in undetectable IFN-γ levels in the supernatant 24 hours after CAR-T-cell addition, confirming effective cytokine neutralization. Importantly, TNF-α secretion was substantially reduced, further supporting the notion that IFN-γ acts as a key upstream mediator of OTOT toxicity by amplifying local inflammatory signals (**Fig. 5D, 5E**).

To further mitigate CD123 CAR-T cell-mediated toxicity, we tested dasatinib, a tyrosine kinase inhibitor previously shown to reversibly suppress CAR-T cell activation at low concentrations by transient inhibition of CAR signaling ^37,38^. In our tissue slice model, dasatinib markedly decreased cytokine secretion when applied at the start of co-culture (D0h) (**Fig. 5D**). Even when added 6 hours after co-culture (D6h), cytokine levels remained significantly lower than in CAR-T-cell controls (**Fig. 5D, 5E**).

To ensure that these interventions did not impair CAR-T-cell function, we assessed cytotoxicity against CD123-expressing CAL1 cells. Both anti-IFN-γ and dasatinib conditions maintained robust CAR-T-cell-mediated killing, indicating preserved cytotoxic activity despite reduced toxicity (**Fig. 5F**). The RPMI control showed no evidence of basal activation, confirming that CAR-T-cell activity was antigen-dependent (**Fig. Sup. 5B**).

By neutralizing IFN-γ or pharmacologically inhibiting CAR-T cell activity, we observed a marked reduction in activation, thereby limiting excessive inflammatory responses without permanently impairing CAR-T cell function. This temporal control may be particularly relevant in contexts where inflammation can further amplify antigen expression in healthy tissues. These results highlight the value of the platform as an experimental model for testing strategies aimed at enhancing the safety of CAR-T therapies.

### Organotypic lung slices guide the selection of safer HER2 CAR-T cells by revealing affinity-dependent OTOT toxicity

We next leveraged the organotypic platform to guide CAR design strategies aimed at reducing OTOT toxicity. Considering the critical role of CAR antigen-binding affinity and architecture, we applied this framework to HER2-targeting CAR-T cells, including HER2 low-affinity second-generation, HER2 high-affinity second-generation, and HER2 high-affinity third-generation constructs in healthy human lung tissue ^9,39^ (**Fig Sup 6A, 6B, 2A**).

Before the organotypic experiments, we first confirmed in vitro the differential activation potential of low- and high-affinity HER2 CAR-T cells by measuring calcium signaling in response to moderate HER2 expression in triple-negative breast cancer cells (MDA-MB-453, reflecting healthy tissue levels) and high HER2 expression in a HER2+ breast cancer cell line (SKBR3) (**Fig. 6, Sup. 6C**). Quantification showed that both low- and high-affinity CARs were strongly activated by high HER2-expressing cells (**Fig. 6B, 6C top**), whereas only high-affinity CARs responded to moderate expression, with no difference between 2nd- and 3rd-generation constructs (**Fig. 6B, 6C bottom**).

**Fig. 6:**
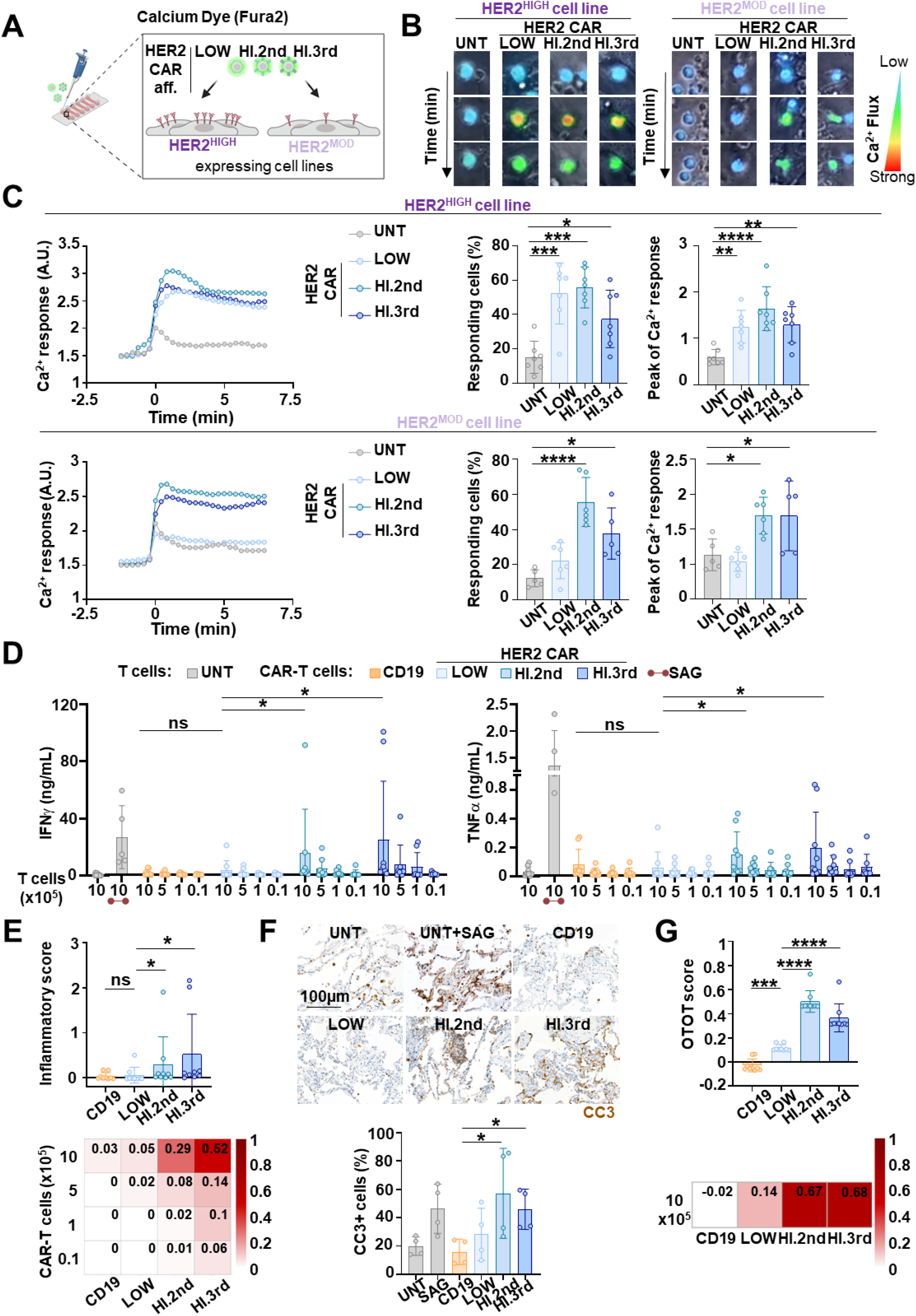
HER2 low-affinity CAR-T cells exhibit reduced OTOT in a human lung organotypic model. **A)** Schematic of the calcium flux assay. HER2 CAR-T cells with low affinity (LOW), high affinity second-generation (HI.2nd), and high affinity third-generation (HI.3rd) and UNT T cells were loaded with Fura-2 and co-cultured with cell lines expressing high or moderate levels of HER2. Calcium activity was recorded every 30s over 10min by live-cell imaging. **B)** Representative time-lapse images of calcium flux in CAR-T cells interacting with HER2^high^ and HER^mod^ cell lines. Ratiometric fluorescence change from blue to yellow/red (strong response) and green (moderate response). **C)** Calcium response kinetics, percentage of responding cells, and peak calcium responses (Δ maximum vs baseline) following co-culture with HER2^high^ and HER^mod^ cell lines (n = 4 - 7, replicates). **D)** IFN-γ and TNF-α concentrations were measured after 24 hours of co-culture with lung tissue and are presented in bar graphs. Different CAR/UNT ratios were used, as described in Figure 1. UNT alone = negative control; 1 µg/ml superantigen with UNT (SAG) = positive control (red underline). Data are mean values n = 4 tissue donors; n = 3 T-cell donors; duplicate slices. **E)** Inflammatory score calculated from IFN-γ and TNF-α secretion. Bar graph shows 10×10⁵ CAR-T cells per slice; heatmap summarizes all tested doses. Scores were normalized from 0 (UNT) to 1 (SAG). **F)** Images showing CC3 (brown) detected by IHC for the condition with 10×10^5^ cells. Quantification of the % of CC3+ cells is presented (n = 6 tissue sections per condition). **G)** OTOT score combining IFN-γ, TNF-α, and CC3+ cell for UNT and CAR-T cells at 10×10^5^ cells per lung slice. Scores were normalized between 0 (UNT) and 1 (SAG). Statistical significance was determined using a two-tailed Mann-Whitney test or a one-way ANOVA with Tukey’s multiple comparison test; *, p<0.05; **, p<0.01; ***, p<0.001**; ****,** p<0.0001.

We then assessed whether this reduced sensitivity would translate into lower toxicity in the organotypic lung tissue model. CD19 CAR-T-cells were included as a negative control. Levels of IFN-γ and TNF-α produced by CD19 CAR and low-affinity HER2 CAR-T-cells were significantly lower compared to those produced by high-affinity CAR-T-cells. These findings suggest that the use of low-affinity HER2 CARs may offer a safer therapeutic profile than high-affinity CARs (**Fig. 6D, 6E**).

Tissue damage analysis via CC3 staining showed that high-affinity HER2 CAR-T cells induced significantly more apoptosis, whereas low-affinity CARs induced CC3 levels comparable to CD19 CAR-T cells, indicating minimal cytotoxicity in healthy lung tissue (**Fig. 6F, 6G**). Both low- and high-affinity CAR-T cells retained comparable cytokine secretion against HER2-high tumor cell lines, with no basal activation, confirming that reduced affinity does not impair activity against high-antigen-expressing tumor targets (**Fig. Sup. 6D**). Organotypic lung slices guide safer HER2 CAR-T cells by revealing affinity-dependent OTOT toxicity. Collectively, these results demonstrate the discriminative power of our model and establish its value as a preclinical framework for rational CAR design, balancing potent antitumor activity with reduced risk of OTOT toxicity in healthy tissues.

## Discussion

Predicting OTOT toxicity in CAR-T therapy remains challenging, largely due to the limited predictive value of current preclinical models and of antigen-expression-based risk assessment alone ^7,20^. To address this gap, we developed an *ex vivo* organotypic slice assay that advances CAR-T cell evaluation by (1) providing a physiologically relevant platform to assess efficacy and safety while capturing both basal and tissue-specific inflammation and (2) enabling functional evaluation of OTOT toxicity to capture target-specific toxicity profiles and guide rational CAR-T cell design.

CAR constructs targeting HER2, EGFR, and MSLN induced measurable tissue damage in healthy lung slices, confirming their potential for pulmonary toxicity and demonstrating the assay’s ability to capture such OTOT effects ^2–4^. IFN-γ and TNF-α secretion strongly correlated with CC3 levels and CAR-T cell activation, establishing cytokine profiling as a rapid and reliable surrogate for cytotoxicity. A major observation was that low antigen detection by IHC did not preclude robust CAR-T cell activation and tissue damage. This finding reinforces the notion that antigen expression, although essential for target selection, does not fully predict the functional consequences of CAR-T cell engagement with normal tissues.

Cytokine secretion levels in tumor tissue slices remained comparable to those in healthy tissues despite higher target antigen expression. This was particularly evident in some tumor slices in which HER2 or MSLN expression was increased relative to non-tumoral lung tissue, without a proportional increase in CAR-T cell activation. These observations suggest that the tumor microenvironment may impose functional constraints on CAR-T cell responses, even when target antigen is detectable. These constraints may involve antigen accessibility, stromal organization, immune suppression, local cytokine networks, and intrinsic resistance mechanisms. This is consistent with previous studies showing that tumor-derived tissue models retain tumor-associated features that regulate immune responses independently of antigen expression ^23,44^. Conversely, healthy tissue slices may also exhibit context-dependent features that influence CAR-T cell responses. Since these slices were derived from tumor-adjacent tissue, they may display baseline inflammatory states, while tissue processing can further induce local stress responses ^45,46^. To capture such environment-driven changes, healthy tissue slices were preconditioned with rIFN-γ, revealing a target-specific increase in CD123 CAR-T cell-mediated toxicity. Notably, this inflammatory priming did not similarly enhance toxicity across EGFR-, CD19-, MSLN-, or HER2-targeted CAR-T cells, further demonstrating that inflammatory signals modulate CAR-T cell activity in a target-dependent manner rather than acting as a universal amplifier of toxicity. These findings demonstrate that different targets exhibit distinct functional responses and reinforce the need for functional models.

The predictive capacity of the platform was further supported using ROR1 CAR-T cells, which have been associated with OTOT safety concerns in preclinical models due to low-level expression of ROR1 in lung compartments, where CAR-T cell engagement and infiltration have been reported ^13,27^. In our study, we found functional concordance between the *in vivo* and *ex vivo* organotypic models, with both systems showing a dose-dependent increase in IFN-γ secretion in response to ROR1 CAR-T cells. Consistently, dose-dependent toxicity was observed across all CAR-T cell constructs, in line with clinical reports of pulmonary toxicity associated with higher infusion doses of HER2- or MSLN-targeting CAR-T cells, highlighting the critical role of cell dose in determining toxicity severity ^9,10^. These findings emphasize that OTOT toxicity represents not only a safety concern but also a limitation to CAR-T cell dosing and therapeutic efficacy. Strategies that reduce normal tissue toxicity while preserving antitumor activity may therefore expand the therapeutic window by enabling administration of higher CAR-T cell doses.

It is important to note that while overall ROR1 CAR-T cell density appears broadly comparable between *ex vivo* and *in vivo* contexts, their spatial distribution differs: intravenous delivery *in vivo* enables organized trafficking, whereas direct application onto slices produces an artificial distribution. However, a study with CDH17-targeting CAR-T cells on human healthy colon slices demonstrated that although CDH17 is expressed across the tissue, its localization at cell-cell junctions restricts CAR-T cell access, resulting in minimal toxicity ^21^. Thus, despite differences in spatial distribution, the assay captures key determinants of CAR-T cell-mediated tissue damage, including target accessibility and tissue organization. Indeed, epithelial and stromal cell-associated antigens, such as EGFR, HER2, and MSLN, are often clustered on continuous tissue surfaces, which may facilitate repeated CAR-T cell engagements. In contrast, dispersed targets like CD19, may provide fewer opportunities for sustained interactions, potentially limiting inflammatory loops. Together, these findings support that target display distinct functional behaviors determined by antigen localization and tissue organization.

In addition, the platform also captures organ-specific tissue contexts. TNC-targeting CAR-T cells elicited robust cytokine release and increased apoptotic cell death across multiple cynomolgus macaque tissues, while most other CAR-T cell constructs remained largely inactive. Notably, BCAN, CSPG4, and PTPRZ1 are predominantly restricted to the brain and central nervous system, whereas TNC, as a component of the extracellular matrix, shows distribution across multiple tissues ^51–56^. However, TNC expression could also be induced by the mechanical stress or inflammation associated with organotypic slice preparation ^31^. The observed activity may be enhanced by inflammation-driven TNC upregulation, or alternatively reflect off-target recognition of cynomolgus macaque antigens by the CAR construct. Importantly, the platform enables the assessment of CAR-T cell activity across a broad range of organ-derived tissue slices, supporting the evaluation of tissue-specific reactivity in diverse physiological contexts and improving preclinical toxicity profiling.

A key objective of this platform is to identify strategies to reduce the risk of OTOT toxicity induced by CAR-T cells. Pharmacological interventions provide an avenue to modulate tissue injury. Dasatinib directly attenuated CAR-T activity while maintaining anti-tumor efficacy, whereas anti-IFN-γ antibodies selectively dampened CD123-associated local inflammatory loops, reducing tissue toxicity without compromising cytotoxic function ^37,38^. Moreover, receptor affinity emerged as a critical determinant of CAR-T cytotoxicity. Low-affinity HER2-targeting CARs induced significantly less tissue damage, consistent with evidence that tuning scFv affinity can modulate sensitivity and discriminate tumor from normal cells^16,57,58^. The platform can also be applied to evaluate next-generation CAR architectures, such as logic-gated CAR-T cells or NK-based constructs ^59–62^.

The organotypic slice assay supports CAR design development and facilitates the identification of pharmacological strategies to mitigate toxicity. Although prospective patient-specific validation would be highly valuable, it is inherently limited by the availability of matched healthy tissues, as tissues collected adjacent to the tumor may not correspond to the organs susceptible to OTOT toxicity. Instead, the primary strength of this platform lies in providing a broadly applicable human tissue-based framework for comparing the safety profiles of CAR-T-cell products across different CAR-target-tissue combinations during preclinical development and for inclusion in regulatory submission packages. This is particularly relevant given that our findings demonstrate that OTOT toxicity is determined by the interaction between the CAR, its target antigen, and the surrounding tissue environment. This platform may further guide CAR-T-cell dosing strategies, patient stratification based on organ vulnerability, and the identification of critical monitoring windows. Together, the organotypic slice assay provides a framework for the rational design of safer and more effective CAR-T therapies.

## Materials and Methods

### Study approval

Human studies were conducted in accordance with French biomedical research regulations and the Declaration of Helsink. Institutional review board approval was granted by CPP Ile de France (approval number 00001072, August 27, 2012) and by the Commission Cantonale d’Ethique de la Recherche sur l’être humain of Geneva (approval numbers 2022-02109 and 2023-01923). Animal experiments were approved by the Animal Experimentation Ethics Committee (CEEA 17-039) and the French Ministry of Research (APAFiS #15076). Peripheral blood samples were obtained from the French Blood Establishment (EFS; agreement 18/EFS/030) following informed consent. All samples were anonymized prior to analysis.

### Human samples

Ten fresh healthy lung tissue samples and three lung adenocarcinoma specimens were obtained from anonymized patients undergoing surgical resection. Tissues were processed 2-24 h after resection and maintained at 4°C until use. Written informed consent was obtained from all patients in accordance with ethical regulations.

### Non-human primate samples

Tissue samples come from the MOGBX1 Tox study conducted on Cynomolgus monkeys by the French CRO Motac Neurosciences on behalf of OGD2 Pharma. Monkey tissue samples were collected from the group control (1 male #583 and 1 female #856), in accordance with institutional and national ethical regulations governing animal research. Experiments were conducted using fresh healthy lung tissue obtained within 2 to 24 hours following surgical collection. All samples were maintained at 4°C during transport and prior to further processing.

### Cell lines

BxPC3, SKBR3, CAPAN-1, A375, Ge518_PTPRZ1_KI, HEK 293 T and Pan02 were cultured in DMEM (Thermo Fisher Scientific; cat. #31966021) supplemented with 10% FBS and 1% penicillin-streptomycin (P/S; Thermo Fisher Scientific; cat. #15140122); MDA-MB-453 HER2-low in DMEM/F12 (Thermo Fisher Scientific; cat. #11320033) with 10% FBS, 1% GlutaMAX (Thermo Fisher Scientific; cat. #35050061), and 1% P/S; NALM6 and CAL-1 in RPMI (Thermo Fisher Scientific; cat. #61870044) with 10% FBS and 1% P/S. The Ge518-PTPRZ1_KI cell line was generated from a patient with glioblastoma and transduced with the α-carbonic anhydrase and fibronectin type III extracellular domains (aa 1-419), followed by the transmembrane domain (aa 1,636-1,671) of human PTPRZ1 ^28^.

### Generation of human and murine CAR-T cells

scFvs (aCD19, HER2, mesothelin), signal peptide, CD8 hinge, CD8 transmembrane regions and intracellular domains from 41BB and CD3Z), were synthesized by Genscript and cloned into the third-generation lentiviral vector pCCL under the control of EF1α promoter ^11,41–43^. EGFR-targeting scFvs were previously described in earlier studies ^42^. Lentiviral vectors were produced after transfection of 293FT and tittered in Jurkat cells^9,39,42^.

#### Generation of human CAR-T cells targeting EGFR, Mesothelin, CD19, and HER2 (Low, High - Second and Third Generation)

CAR-T cells were generated from PBMCs isolated from four healthy donors by Ficoll density gradient centrifugation (800 × g, 20 min, room temperature). T cells were purified using the Pan T Cell Isolation Kit (Miltenyi Biotec; cat. #130-096-535) and cultured in RPMI 1640 (Thermo Fisher Scientific; cat. #21875) supplemented with 10% FBS (Thermo Fisher Scientific; cat. #A3840002) and 1% penicillin-streptomycin (Thermo Fisher Scientific; cat. #15140130). T cells were activated with TransAct (Miltenyi Biotec; cat. #130-111-160) and cultured with recombinant human IL-7 (10 ng/mL; Miltenyi Biotec; cat. #130-095-362) and IL-15 (10 ng/mL; Miltenyi Biotec; cat. #130-095-765). Two days after activation, cells were transduced in 48-well plates with the indicated CAR constructs using construct-specific MOIs (2 for CD19 and EGFR, 2.5 for HER2 LOW, HER2 HI.2nd, and HER2 HI.3rd, and 8 for Mesothelin). Cells were harvested three days later and maintained at 2×x10⁶ cells/mL. After one week, CAR-T cells were either used for functional assays or cryopreserved (Sigma-Aldrich; cat. #C2874-100ML). CAR expression was confirmed by flow cytometry.

#### Generation of human CAR-T cells targeting BCAN, CSPG4, PTPRZ1, and TNC

CAR-T cells were generated from one healthy donor. T cells were purified from a buffy coat using the EasySep™ Direct Human T Cell Isolation Kit (Stemcell Technologies, cat# 19661). After purification, T cells density was adjusted to 1 x 10^6^ cells/mL and activated with anti-CD3/anti-CD28 Dynabeads (Gibco, 11141D) at 1:1 ratio. T cells were incubated for 48h, then beads were magnetically removed and activated T cells were washed with MaxCyte Electroporation buffer (Cytiva Life Sciences, EPB1). mRNA was produced using the MEGAscript™ T7 Transcription Kit (Thermofisher, cat# AM1334) and later cleaned with the MEGAclear Transcription Clean-Up Kit (Invitrogen, cat# AM1908). T cells were electroporated using 3.3 μg of mRNA/10^6^ T cells using ExPERT GTx (MaxCyte). T cells were kept in culture at 37°C with 5% CO_2_ in RPMI 1640 medium (Gibco, cat# 61870) supplemented with 10% fetal bovine serum (Sigma, cat# F7524-500mL), 1 mmol/L sodium pyruvate (Gibco, cat# 11360-070), 10 mmol/L HEPES (Gibco, cat# 15630-056), and 1% penicillin-streptomycin (Gibco, cat# 15140-122). After the first 2 hours of incubation, the medium was supplemented with 30 IU/mL recombinant human interleukin 2 (Proleukin, Roche). Electroporated T cells were kept in culture overnight and then frozen.

#### Generation of human CAR-T cells targeting CD123

CD123 CAR-T cells were produced from PBMCs of healthy donors from the French Blood Establishment (EFS, Bourgogne/Franche-Comté, Besançon), all of whom consented to their blood being used for research purposes ^36^. Briefly, the PBMCs were isolated by Ficoll gradient density centrifugation using Ficoll-Paque (Avantor, Cat# 17-1440). T cells were selected using CD4/ CD8 human microbeads (CD4 MicroBeads, human, Miltenyi Biotec, Cat# 130-097-048; CD8 MicroBeads, human, Miltenyi Biotec, Cat# 130-045-201) and activated using polymeric nanomatrix structure with CD3 and CD28 (T cell TransAct^TM^, human, Miltenyi Biotec, Cat# 130-111-160). T cells cultivated in TexMACS medium (Miltenyi Biotec, Cat# 130-097-196) enriched with 5% heat-inactivated human serum (EFS Bourgogne/Franche-Comté, Besançon), 1% penicillin-streptomycin (Gibco, Cat# 15140122), recombinant human IL-7 (25ng/mL, Miltenyi Biotec, Cat# 130-095-362) and recombinant human IL-15 (25 ng/mL, Miltenyi Biotec, Cat# 130-095-765). Forty-eight hours later, T cells were transduced at a MOI 2 in 24-well plates with 1-hour spinoculation at 800g, 32°C. Every 2 days, T cells were counted and returned to the medium at 1 x 10^6^ T cells/mL. Seven days after transduction, CAR expression was determined by flow cytometry using G4S linker labeling (Cell Signaling Technology, Cat# E7O2V). As control, untransduced activated T cells from the same healthy donor were produced for each experiment ^36^.

#### Generation of murine CAR-T cells targeting ROR1 Retrovirus production

Retroviral supernatant was generated using Plat-E cells. Cells were transiently transfected with a murine ROR1 (R11)-CAR encoding plasmid and the packaging construct pCL-10A1 using the Effectene Transfection Reagent (QIAGEN) according to the manufacturer’s instructions. Briefly, plasmid DNA was mixed with Enhancer and Effectene reagent to generate transfection complexes, which were added to the cells. Viral supernatant was collected 48 and 72h after transfection, pooled, filtered to remove cellular debris, and used for the transduction of murine T cells.

Spleens were harvested from C57BL/6 CD45.1 mice and were mechanically dissociated through a 70 µm cell strainer to obtain single-cell suspensions. Cells were resuspended in red blood cell lysis buffer and incubated for 7 minutes at room temperature to lyse erythrocytes. T cells were isolated using the Miltenyi Mouse T Cell Isolation Kit (Miltenyi Biotec; cat #130-095-130). 24-well plates were coated with 5 µg/mL anti-CD3 antibody diluted in 300 µL PBS and incubated overnight at 4°C. CAR-T cells were generated by transducing activated T cells two days post-activation in complete RPMI medium (10% FCS, 1% penicillin-streptomycin, 50 μM 2-mercaptoethanol in RPMI-1640 medium) supplemented with 50 U/mL IL-2. Retroviral transductions were carried out in the presence of Polybrene (1:1000; Merk; #TR1003G), followed by spinoculation at 800xg for 90 minutes at 32°C. Cells were incubated for 3-4 hours at 37°C. After incubation, the viral supernatant was removed, and fresh complete RPMI medium was added. On day 3, half of the culture medium was replaced with fresh medium containing IL-2 (50 U/mL). On day 4, cells were centrifuged, transferred to new plates, and cultured in medium supplemented with IL-2, IL-7, and IL-15 (Miltenyi Biotec; cat #130-120-662; #130-094-066; #130-094-072). Transduction efficiency was assessed by flow cytometry using transduction markers (mCD19; Biolegend; cat #152408). On day 7, the CAR-T cells were used for functional assays.

### Organotypic slice assay

Tissue slices were prepared as previously described ^26^. Human healthy lung, lung tumor, and mouse tumor tissues were embedded in 5% low-melting agarose (Sigma-Aldrich; cat. #A0701-25G) in PBS (Thermo Fisher Scientific; cat. #10010001) and sectioned at 400 µm using a Leica VT1200S vibratome (RRID:SCR_018453) in ice-cold PBS. Slices were placed individually into culture inserts within 24-well plates (Merk; cat #PITP01250) and a stainless-steel washer (4 mm inner diameter) was positioned over each slice to confine cell application. total of 1×x10⁶ T cells were added per slice at CAR-T/UNT ratios of 10/0, 5/5, 1/9, or 0.1/9.9 (×x10⁵), maintaining a constant total cell number. After 30-60 min at 37°C, 5% CO₂, complete medium (400 µL) was added, washers were removed, and superantigen was added where indicated (1:1000). Co-cultures were maintained for 24 h. Cytokines in supernatants were quantified using the LEGENDplex assay (BioLegend) and acquired on an Attune NxT cytometer (Thermo Fisher Scientific). Tissue slices were then fixed in 10% neutral buffered formalin (Sigma-Aldrich; cat. #HT501128), paraffin-embedded, and stained for CC3 using anti-CC3 (Ozyme; cat. #CST9661; 1:400), HRP-conjugated secondary antibody (Abcam; cat. #ab64264), and DAB. LEGENDplex data were analyzed with Qognit and IHC images with QuPath. For flow cytometry, T cells were pre-labeled with CFSE (1:5000), co-cultured for 24 h, and tissue were dissociated into single-cell suspensions before analysis.

To quantify the inflammatory response, cytokine levels were normalized to the negative (UNT) and positive (SAG) controls to generate an inflammatory score, where UNT = 0 and SAG = 1, using the following formula:

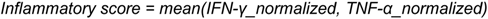

The inflammatory score was then obtained as the mean of normalized IFN-γ and TNF-α levels:

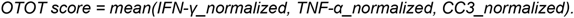

To assess combined inflammatory and cytotoxic effects, an OTOT score was computed using the same normalization for IFN-γ, TNF-α, and CC3:

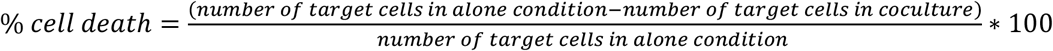

### Spatial transcriptomic

Spatial transcriptomics was performed on formalin-fixed, paraffin-embedded (FFPE) lung tissue sections and performed using the Visium HD platform (10x Genomics), incorporating three specific probes targeting CAR-T cells during the hybridization step to enable their detection. Sequencing was executed on an Illumina NovaSeq 6000 system using an SP flow cell in paired-end mode (200 cycles) with a 5% PhiX spike-in. Data processing was conducted using the standard SpaceRanger pipeline, with customized reference files (genome/transcriptome) to ensure the capture and identification of the added CAR-T specific probes. Analysis was performed within the SpatialX environment (BioTuring) using 8 µm binned data, filtered to retain only bins with a minimum of 10 genes. Initial bins annotation was conducted using the MetaReference model (trained on human tissues) and subsequently refined manually, guided by established cellular types and subtypes to ensure biological accuracy ^43^. CAR-T cells were specifically annotated using CAR-T probes detection, and remaining unassigned bins were removed. Continuous biological signal scores like HER2+, apoptosis or T activation have been derived by bin using AUCell algorithm and transcriptomic signatures.

### Intracellular calcium (Ca^2+^) measurement

Intracellular Ca²⁺ was measured in untransduced (UNT) and CAR-T cells loaded with 1 µM Fura-2 AM (Thermo Fisher Scientific; cat. #F1221) for 30 min at 37°C, 5% CO₂. After washing, 0.5 × 10⁶ T cells were added to HER2-low MDA-MB-453 or HER2-high SKBR3 cells cultured in six-channel IBIDI µ-Slides. Fluorescence images were acquired every 10 s for 10 min at 340 and 380 nm using MetaFluor software. Intracellular Ca²⁺ levels were calculated as the F340/F380 fluorescence ratio. CAR-T cells were considered responsive when the Ca²⁺ signal was at least twofold above background. Data were analyzed using Fiji software.

### *In vitro* cytotoxic functionality

To evaluate CD123 CAR-T mediated cytotoxicity, untransduced or CD123 CAR-T cells were previously labelled using a Cell Proliferation Dye eFluor 450 solution (Invitrogen, Cat# 65-0853) before being co-cultured with CAL-1 cells with effector: target of 1: 1 during 24 hours with or without anti-IFN-γ (10µg/mL, BioXCell, Cat# BE0235) or Dasatinib (50nM, Sigma Aldrich, Cat# BMS-354825). Cell death was assessed by flow cytometry using CD123 (Sony Biotechnology, Cat# 2130050) and 7-AAD labelling (Sony Biotechnology, Cat# 2702020) for cell line detection, and T cells were targeted by fixable viability Dye eFluor labelling. Cell death was calculated using the following formula:

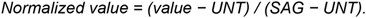

Flow cytometry analysis was performed using a BD Symphony A1 cytometer (BD Biosciences). Data were analysed using FACS Diva software (BD Biosciences).

### Animal Experiment

C57BL/6-CD45.1 and C57BL/6-CD45.2 (C57BL/6J) mice (Charles River Laboratories) were housed at the Cochin Institute Animal Facility. C57BL/6-CD45.2 mice were subcutaneously injected with 1 × 10⁶ PAN02 cells in 100 µL PBS. Seven days later, mice received intravenous ROR1 CD45.1 CAR-T cells (1×x10⁶ or 1×x10⁷), UNT cells (1×x10⁷), or no treatment. Three days later, lungs were collected, sectioned into 400 µm slices as described above, cultured for 24h, and supernatants were collected for cytokine analysis. Parallel lung samples (5 mm) were fixed in 4% paraformaldehyde (PFA), vibratome-sectioned (400 µm), and processed for immunofluorescence.

### Immunofluorescence

Lung slices were stained by immunofluorescence for CD45.1 (to detect CAR-T cells; Biolegend; cat#110714), GP38 (Biolegend; cat#127408) and ROR1 (ThermoFisher Scientific; cat #PA514726) with an appropriate secondary antibody used for ROR1 detection (ThermoFisher Scientific; cat #31635). Imaging was performed using a Confocal Xplorer microscope (Leica Microsystems). Data were analyzed using Fiji software.

### Flow Cytometry

Lungs were enzymatically dissociated into single-cell suspensions using Liberase TM (Merk; cat #5401119001) and DNase I (Merk; cat #4716728001), followed by mechanical disruption. After filtration through a 70 µm strainer, cells were stained with antibodies against EGFR, HER2, CD19, and mesothelin (Biolegend; cat #352904; #324406; #982402; Miltenyi Biotec; cat #130118168), as well as with the viability dye Live/Dead Aqua (Invitrogen; cat #L34966). CAR-T cell expression was measured using a Human Fab-specific antibody (Jackson ImmunoResearch; cat #109-606-006). CAR-T cell activation was assessed using CellTrace™ CFSE (ThermoFisher #C34554. CD3, CD69, CD25, PD1, 4-1BB (Biolegend; cat #317344 #310912 #302636 #329908 #300804)

Flow cytometry analysis was performed using a BD LSRFortessa cytometer (BD Biosciences). Data were analyzed using FlowJo software.

### Immunohistochemistry

Formalin-fixed, paraffin-embedded lung tissue sections were deparaffinized, rehydrated, and subjected to heat-induced epitope retrieval in citrate buffer. Endogenous peroxidase activity was blocked using hydrogen peroxide. Sections were then incubated with primary antibodies against mesothelin (Abcam; cat #ab196235), EGFR (Abcam; cat #ab30), CD19 (Abcam; cat #ab134114), and HER2 (Abcam; cat #ab16901). Detection was performed using an HRP-conjugated secondary antibody and developed with DAB substrate. Data were analyzed using QuPath software.

### RT-qPCR

Total RNA was extracted from lung tissue slices (Macherey-Nagel; cat #740984.50) and reverse-transcribed into cDNA using a reverse transcription kit (ThermoFisher; cat #4368814). qPCR was performed using SYBR Green (ThermoFisher; cat # A25742). CD123 expression was quantified using specific primers and normalized to GAPDH as housekeeping gene. Relative expression was calculated using the 2^-ΔΔCt method with NT samples as calibrator.

## Statistical analysis

Statistical analyses were performed using GraphPad Prism software. Data are presented as mean ± SD. *, p<0.05; **, p<0.01; ***, p<0.001; ****, p<0.0001.

## Acknowledgements

This work was supported by the Ligue contre le Cancer (équipe labellisée), the Innovative Medicines Initiative 2 Joint Undertaking under grant agreement No 116026 (this Joint Undertaking receives support from the European Union’s Horizon 2020 research and innovation program and EFPIA) and ERA-NET TRANSCAN-3 (EC co-funded call 2021, SmartCAR-T). This work was also supported by the ISREC foundation and the Private Foundation of the Geneva University Hospitals (DM). We thank all members of our laboratory for their valuable scientific input and support and OGD2 Pharma for access to the non-human primate samples. We also thank the HiSTIM platform at the Institut Cochin for their technical assistance.

## Authors contributions

Writing-original draft: AM, GB, MF, ED. Conceptualization: AM, VD, FGO, BL, ML, SG, DM, ED. Investigation: AM, GB, IR, IF, SC, BM, MF, KZM, DMB, MF, LV, SD, FM, AT, MC, XF. Methodology: AM, GB, IR, MF Resources: ALM, BB, MP, DD, LDJM. Funding acquisition: ED Data curation: AM Validation: AM, ED Supervision: ED Formal analysis: AM Project administration: AM, ED Visualization: AM, FM

---

**Supplementary Fig. 1:**
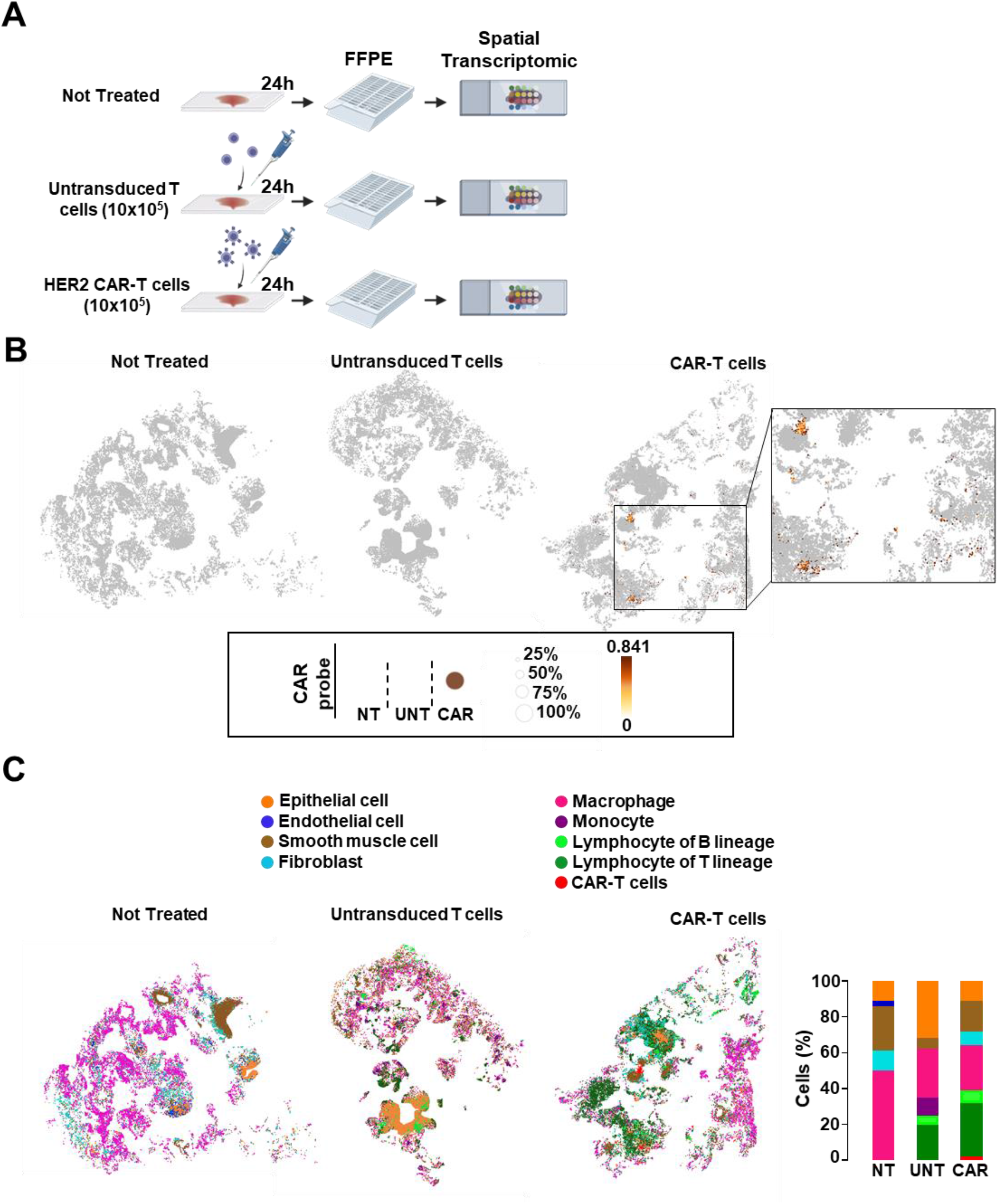
Organotypic human lung slices preserve native tissue architecture and cellular diversity. **A)** Schematic of the experimental design for spatial transcriptomics. Three lung slices were used: untreated control (NT), untransduced T cells (UNT), and HER2 CAR-T-treated (1×x10⁶ cells). After 24 hours of co-culture, tissues were fixed, embedded in FFPE, and processed for spatial transcriptomic analysis. **B)** Spatial map of the three conditions (NT, UNT, and HER2 CAR-T). Each dot represents a capture spot, with signal intensity corresponding to the expression level of the CAR probe, with corresponding quantification shown. **C)** Spatial maps across the three conditions, highlighting the spatial organization of distinct cellular populations. Quantifications are presented in bar graphs.

**Supplementary Fig. 2:**
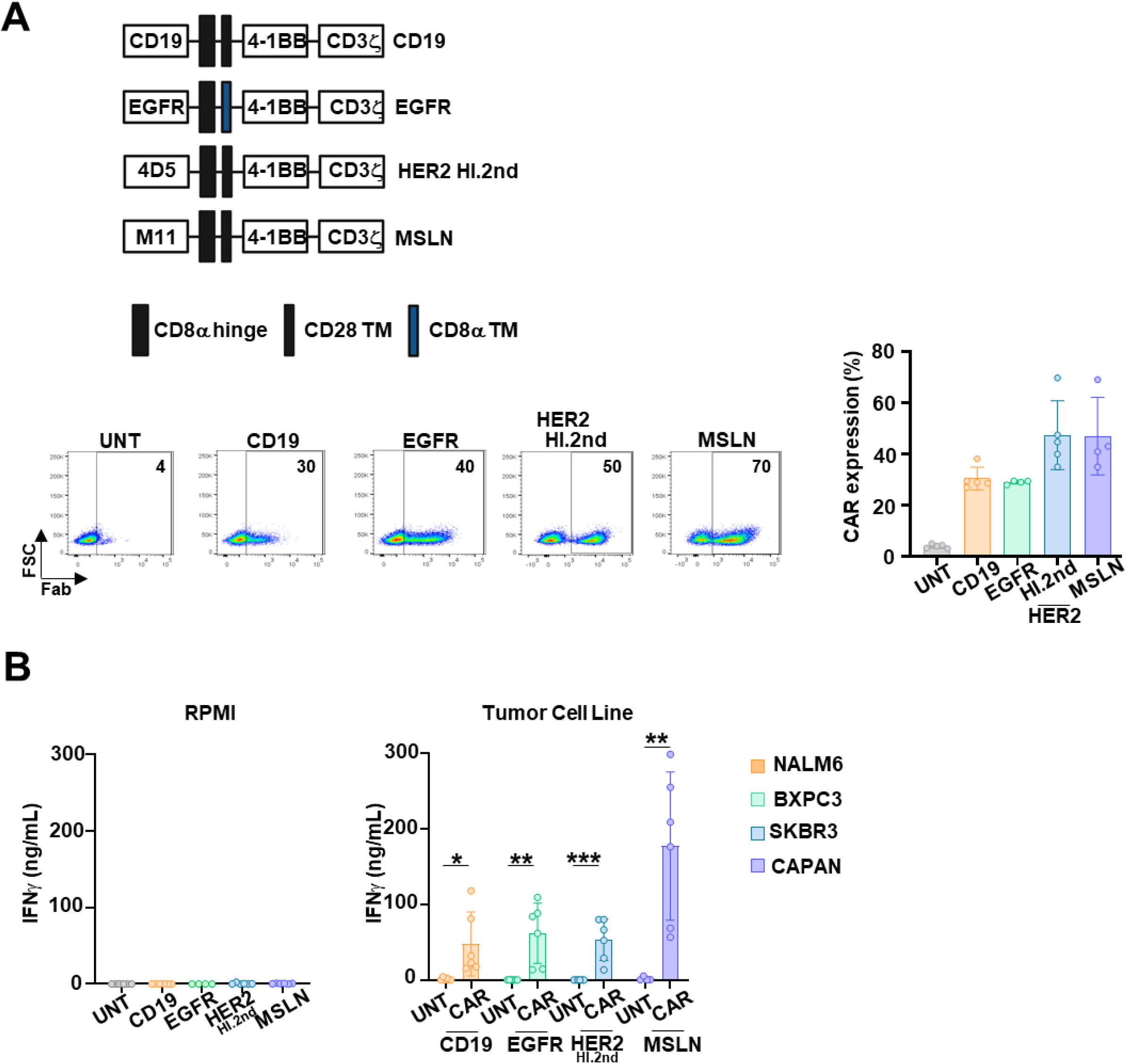
Characterization of CAR expression and *in vitro* functionality of CAR-T cells. **A)** Schematic representation of the CD19, EGFR, HER2 high-affinity, and MSLN CAR constructs, showing the scFv, hinge, transmembrane, and intracellular signaling domains (top). Flow cytometry dot plot showing CAR surface expression, with associated quantification shown below (bottom) (means ± SD). **B)** IFN-γ concentrations produced by CD19, EGFR, HER2 high-affinity, and MSLN CAR-T cells after 24 hours of culture in RPMI medium alone (left) or after co-culture with their respective target cell lines: NALM6 (CD19 CAR-T), BXPC3 (EGFR CAR-T), SKBR3 (HER2 high-affinity CAR-T), and CAPAN (MSLN CAR-T); at an effector-to-target (E:T) ratio of 1:1 (right) (means ± SD) (n=3, experiments performed in duplicate). Statistical significance was determined using a two-tailed unpaired t-test; *, p < 0.05; **, p < 0.01; ***, p < 0.001; **, p < 0.0001.

**Supplementary Fig. 3:**
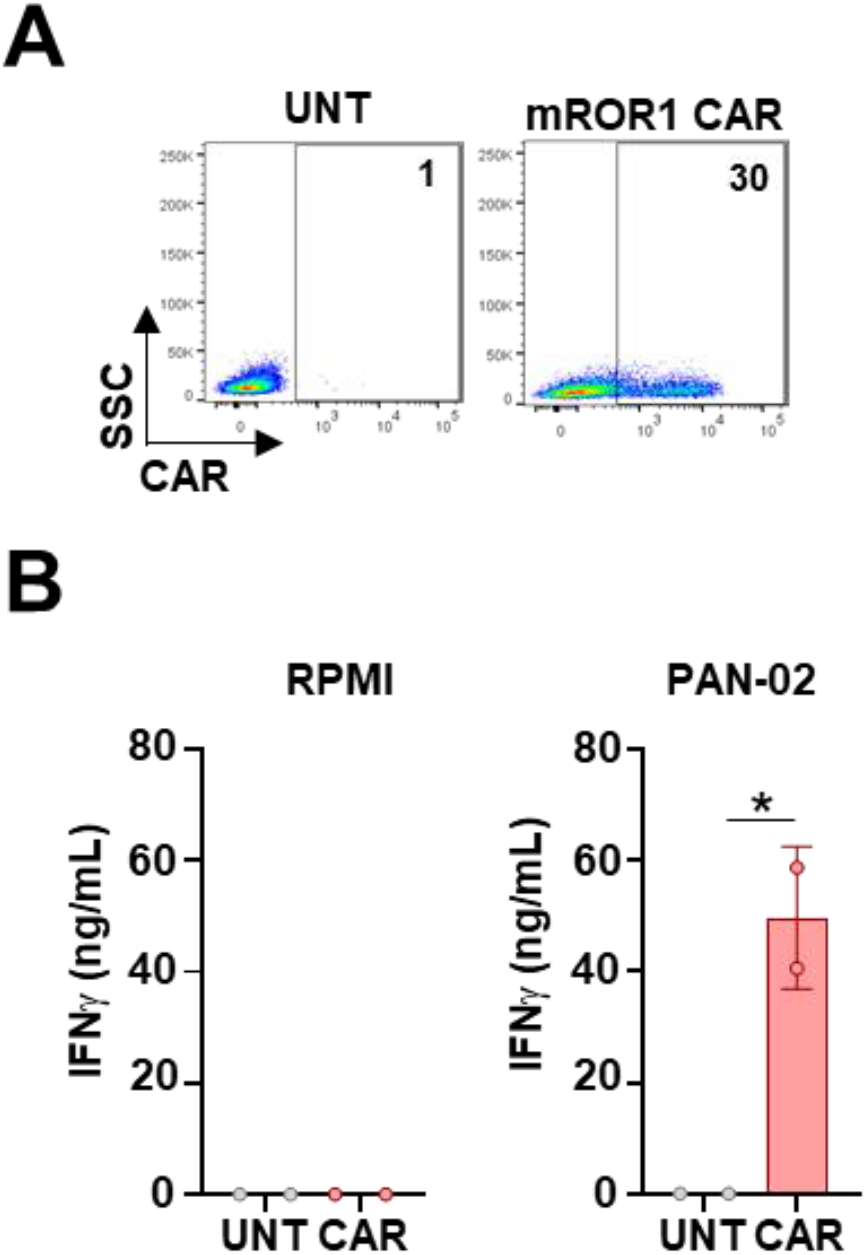
Characterization of CAR expression and *in vitro* functionality of ROR1 murine CAR-T cells. **A)** Flow cytometry analysis of CAR expression on murine cells after retroviral transduction with the ROR1 CAR construct. CAR expression is shown. **B)** IFN-γ concentrations produced by ROR1 CAR-T cells after 24 hours of culture in RPMI medium alone (left) or following co-culture with their ROR1+ target cell line PAN02 (right), shown as mean ± SD. Statistical significance was determined using the Mann-Whitney test; *, p < 0.05.

**Supplementary Fig. 4:**
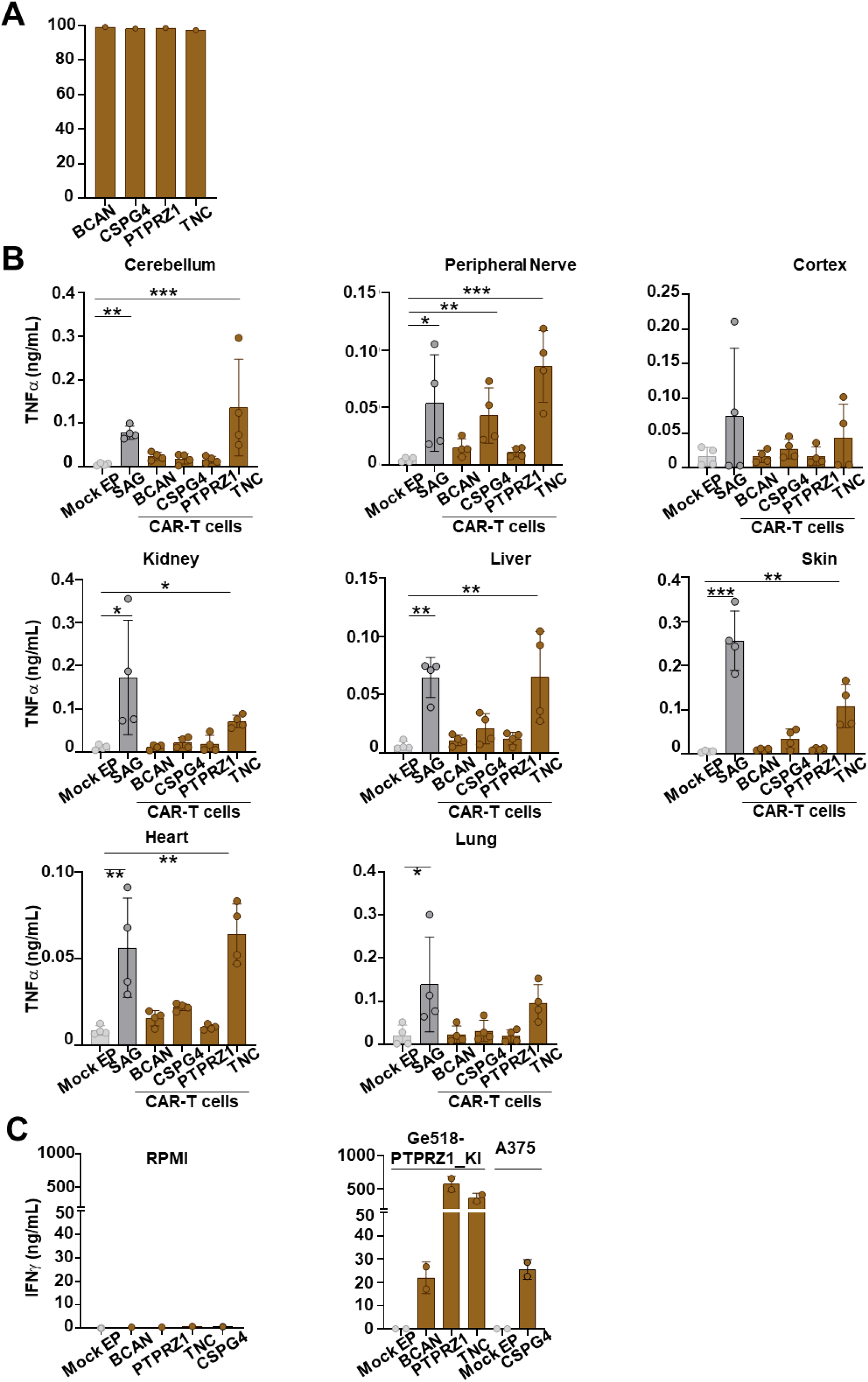
Cytokine secretion by glioblastoma-targeting CAR-T cells across healthy tissues and target cell lines. **A)** Quantification of CAR expression levels. **B)** Bar graph showing TNF-α secretion by the different CAR-T cells (1×x10^6^ cells per slice) across various healthy tissues. Data represent n = 2 cynomolgus macaques, performed in duplicate (mean ± SD). **C)** IFN-γ production by CAR-T cells after 24 hours of culture either in RPMI medium alone (left) or following co-culture with their respective target-expressing cell lines: Ge518-PTPRZ1_KI cell line for BCAN, PTPRZ1 and TNC CAR-T cells and A375 cell line for CSPG4 CAR-T cells and, at an effector-to-target (E:T) ratio of 1:1. Data are presented as mean ± SD. Significance levels were indicated as follows; *, p < 0.05; **, p < 0.01; ***, p < 0.001; ****, p < 0.0001.

**Supplementary Fig. 5:**
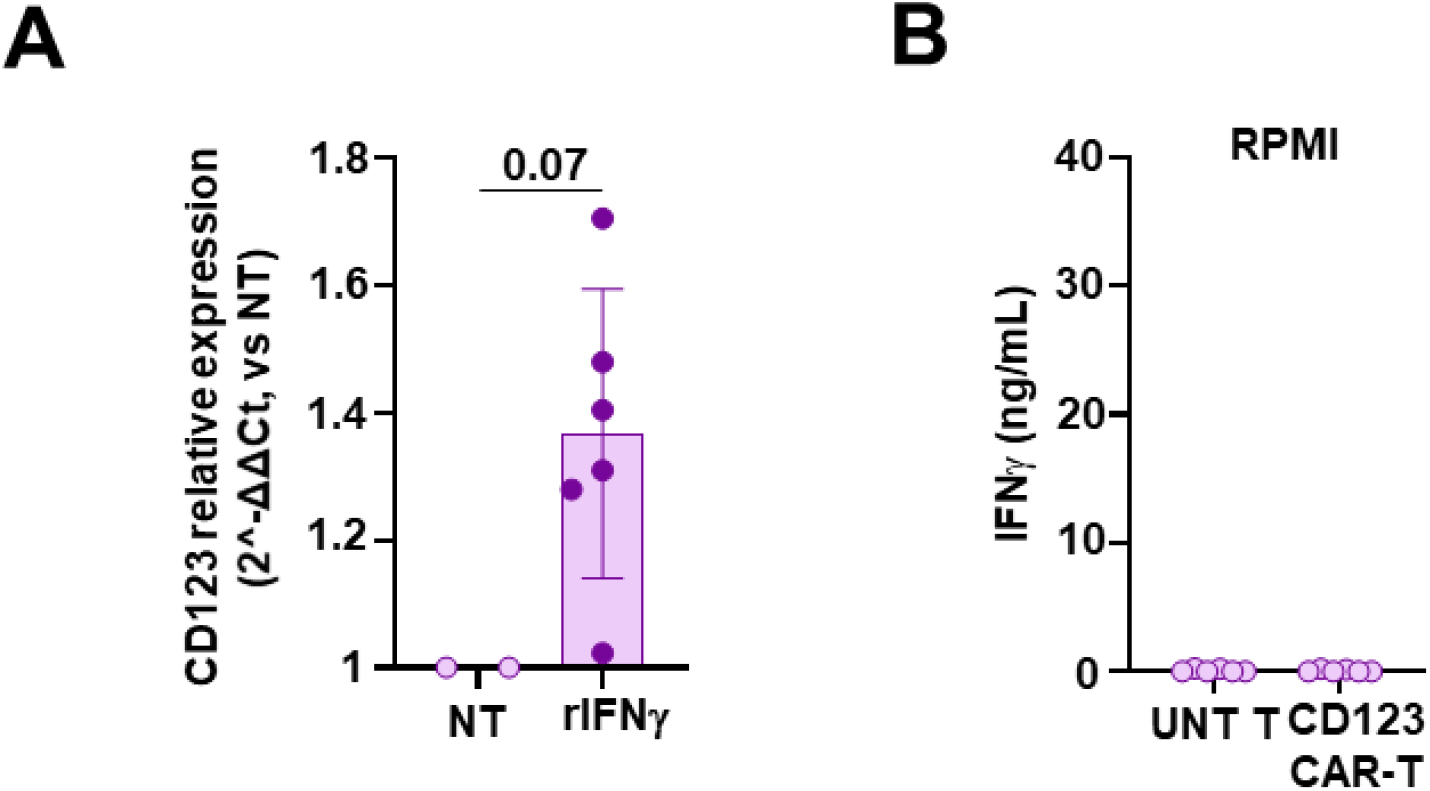
CD123 expression in lung tissue slices and basal cytokine secretion by CAR-T cells. **A)** Bar graphs showing CD123 mRNA expression in lung tissue slices measured by RT-qPCR following 24 h treatment with rIFN-γ (10 pg/mL) or NT control. Data are normalized to NT and presented as relative expression (2^-ΔΔCt). Data represent mean ± SD (n = 2, in triplicate) **B)** IFN-γ concentrations produced by CD123 CAR-T cells after 24 hours of culture in RPMI medium alone. Statistical significance was assessed using an unpaired t-test

**Supplementary Fig. 6:**
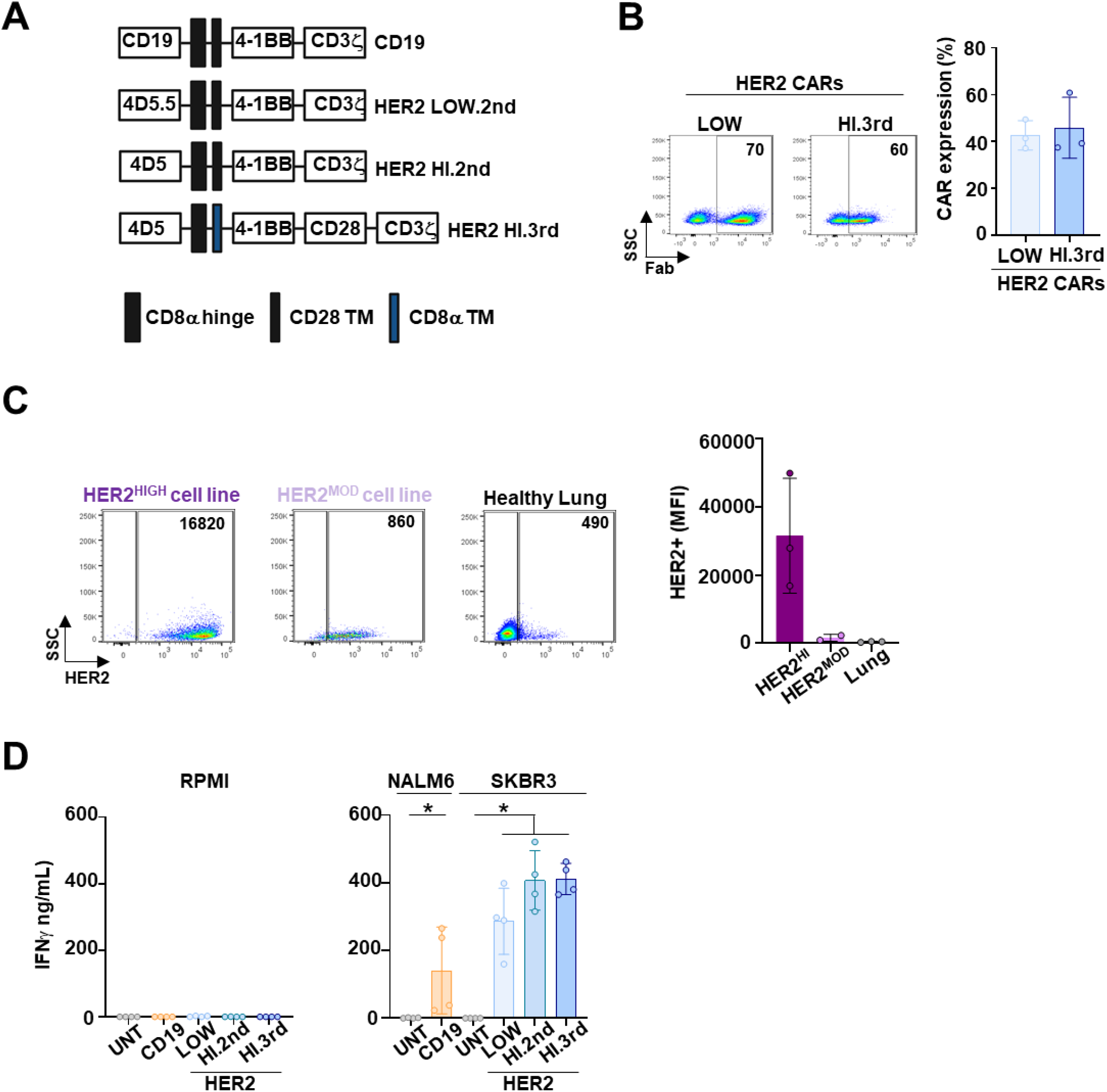
Functional characterization of UNT and CAR-T cells with varying HER2 affinities. **A)** Schematic of CD19, HER2 low-affinity (LOW), HER2 high-affinity second-generation (HI.2nd), and HER2 high-affinity third-generation (HI.3rd) CAR constructs. **B)** Flow cytometry analysis and quantification of CAR surface expression. **C)** Flow cytometry analysis showing HER2 expression in cell lines with high (SKBR3) and moderate (MDA-MB-453 HER2-low) HER2 expression and in lung tissue. The associated quantification shows the mean fluorescence intensity of HER2+ cells. **D)** FN-γ secretion by UNT, CD19, HER2 LOW, HER2 HI.2nd, and HER2 HI.3rd CAR-T cells cultured alone or with their respective target cell lines (NALM6 for CD19; SKBR3 for HER2) for 24 h (n = 4). Statistical significance was determined using a two-tailed Mann-Whitney test; *P < 0.05.

## Notes

### Competing Interest Statement

The authors have declared no competing interest.

